# Large home range scavengers support higher rates of carcass removal

**DOI:** 10.1101/2020.02.07.938415

**Authors:** Cayetano Gutiérrez-Cánovas, Marcos Moleón, Patricia Mateo-Tomás, Pedro P. Olea, Esther Sebastián-González, José Antonio Sánchez-Zapata

**Affiliations:** Freshwater Ecology, Hydrology and Management (FEHM) lab, Departament de Biologia Evolutiva, Ecologia i Ciències Ambientals, Facultat de Biologia, Universitat de Barcelona, Diagonal, 643, 08028 Barcelona, Spain; Centre of Molecular and Environmental Biology (CBMA), Department of Biology, University of Minho, Campus of Gualtar, 4710-057 Braga, Portugal; Institute of Science and Innovation for Bio-Sustainability (IB-S), University of Minho, Campus of Gualtar, 4710-057 Braga, Portugal; Department of Zoology, University of Granada, Avda. de Fuentenueva, s/n, 18071 Granada, Spain; Research Unit of Biodiversity (UO/CSIC/PA), Oviedo University, 33600 Mieres, Spain; Centre for Functional Ecology, Department of Life Sciences, University of Coimbra, Calçada Martim de Freitas, 3000-456 Coimbra, Portugal; Departamento de Ecología, Universidad Autónoma de Madrid, Madrid, Spain; Centro de Investigación en Biodiversidad y Cambio Global (CIBC-UAM), Universidad Autónoma de Madrid, Madrid, Spain; Department of Applied Biology, Miguel Hernández University. Avda. Universidad s/n 03202 Elche, Alicante, Spain

**Keywords:** biodiversity - ecosystem functioning, carrion consumption, ecosystem services, functional diversity, functional identity, functional traits, large carnivores, vultures

## Abstract

1. Vertebrate scavenger communities vary in species composition across the globe, and include a wide array of species with diverse ecological strategies and life-histories that support essential ecosystem functions, such as carrion removal. While previous studies have mostly focussed on how community aspects such as species richness and composition affect carrion consumption rates, it remains unclear whether this important function of scavengers is better explained by the dominance of key functional traits or niche complementarity as a result of a diverse functional representation.
2. Here, we test three competitive hypotheses to assess if carrion consumption in vertebrate scavenger communities depends on: i) the presence of key dominant traits (functional identity hypothesis), ii) functional diversity that promotes niche complementarity (functional diversity hypothesis), or iii) the accumulation of individuals and species, irrespective of their trait representation (functional equivalence). To explore these hypotheses, we used five study areas in Spain and South Africa, which represent a gradient of scavenger biodiversity, i.e., ranging from communities dominated by facultative scavengers, such as generalists and meso-predators, to those including vultures and large carnivores.
3. Within study areas, traits that characterise obligate scavengers or large carnivores (e.g. mean home range, proportion of social foragers) were positively linked to rapid carrion consumption, while the biomass of functional groups including facultative scavengers were either weakly or negatively associated with carrion consumption.
4. When combining all study areas, higher rates of carrion consumption were related to scavenger communities dominated by species with large home ranges (e.g. *Gyps* vultures), which was found to be a key trait. In contrast, metrics describing functional diversity (functional dispersion) and functional equivalence (species richness and abundance) had lower predictive power in explaining carrion consumption patterns.
5. Our data support the functional identity hypothesis as a better framework for explaining carrion consumption rates than functional diversity or equivalence. Our findings contribute to understanding the mechanisms sustaining ecosystem functioning in vertebrate communities and reinforce the role of obligate scavengers and large carnivores as keystone species in terrestrial ecosystems.

## Abstract

1. Las comunidades de carroñeros vertebrados, presentes en ecosistemas terrestres de todo el planeta, incluyen una gran variedad de especies con distintas estrategias ecológicas y biológicas que mantienen funciones esenciales en los ecosistemas, como el consumo de carroña. Hasta el momento, la mayoría de los estudios han analizado cómo la riqueza y la composición de especies influyen en el consumo de carroña, desconociéndose si las variaciones en dicho consumo se deben a la dominancia de ciertos rasgos biológicos o a la complementariedad de nichos y rasgos funcionales dentro de las comunidades de carroñeros vertebrados.
2. En este estudio comparamos tres hipótesis alternativas para explicar si los patrones de consumo de carroña en comunidades de carroñeros vertebrados dependen de: i) la presencia de rasgos dominantes clave para esta función (hipótesis de la identidad funcional), ii) la diversidad funcional que promueve la complementariedad de nichos entre especies, iii) la acumulación de individuos y especies, independientemente de sus rasgos funcionales (equivalencia funcional). Para ello, usamos cinco áreas de estudio de España y Sudáfrica que representan un gradiente de biodiversidad de carroñeros, i.e. desde comunidades dominadas por carroñeros facultativos, como generalistas y depredadores de pequeño tamaño, a comunidades con buitres y grandes carnívoros.
3. Dentro de cada área de estudio, nuestros resultados muestran que un mayor consumo de carroña está asociado a rasgos biológicos como áreas de campeo grandes y alta proporción de carroñeros sociales, características de carroñeros obligados (buitres) y grandes carnívoros. Por el contrario, la biomasa de los grupos funcionales que incluyen carroñeros facultativos se asocia de forma débil o incluso negativa con las tasas de consumo de carroña.
4. Analizando todas las áreas de estudio conjuntamente, observamos que las comunidades de carroñeros con mayores tasas de consumo de carroña cuentan con especies con un rasgo funcional dominante: grandes áreas de campeo (e.g. buitres del género *Gyps*). Por el contrario, las métricas de diversidad funcional (dispersión funcional) y equivalencia funcional (riqueza y abundancia de especies) tuvieron una capacidad menor para explicar el consumo de carroña.
5. Nuestro estudio indica que la hipótesis de la identidad funcional explica mejor los cambios en las tasas de consumo de carroña por comunidades de vertebrados que la diversidad o la equivalencia funcional. Estos resultados contribuyen a entender mejor los mecanismos que sustentan la función ecológica de los vertebrados carroñeros, destacando de nuevo la relevancia de los carroñeros obligados y los grandes carnívoros.

## Introduction

Vertebrate scavenger communities include a wide array of species with different ecology and life-history traits (Devault, Rhodes, Shivik, & Ewjtt, 2003; Mateo-Tomás et al., 2015). In terrestrial ecosystems, the scavenger spectrum ranges from vultures, which show specialised morphological, physiological and behavioural adaptations for an obligate scavenging lifestyle (Cortés-Avizanda, Jovani, Donázar, & Grimm, 2014; Moleón, Sánchez-Zapata, Selva, Donázar, & Owen-Smith, 2014; Ruxton & Houston, 2004), to many other species that eat carrion opportunistically. These facultative scavengers include large mammalian carnivores, meso-predators such as birds of prey and meso-carnivores, and omnivorous species such as corvids and suids (Devault et al., 2003; Mateo-Tomás et al., 2015; Pereira, Owen-Smith, & Moleón, 2014). Through carrion consumption, this complex scavenger community supports key ecosystem functions such as nutrient and organic matter cycling and food web stability (Barton et al., 2013; Beasley, Olson, Selva, & DeVault, 2019; Moleón et al., 2014). However, not all scavengers are equally efficient in supporting such functions (Mateo-Tomás, Olea, Moleón, Selva, & Sánchez-Zapata, 2017). Depending on their functional traits, scavengers may contribute differently to the amount of carrion consumed by the community due to different abilities to find carrion (encounter rates) and ingest it (handling time) (Kane, Healy, Guillerme, Ruxton, & Jackson, 2017). Thus, the extirpation of certain species or groups could jeopardise the ecological role of scavenging assemblages (Inger, Per, Cox, & Gaston, 2016; Markandya et al., 2008; Morales-Reyes et al., 2017). However, despite the previous works suggesting that scavenger traits could be key in explaining carrion consumption, no explicit exploration has been performed to date in terrestrial ecosystems.

Although biodiversity has been recognised as a key driver of ecosystem functioning and services (Duncan, Thompson, & Pettorelli, 2015; Hooper et al., 2012), there are still many uncertainties about which community features and ecological mechanisms best predict carrion consumption in terrestrial ecosystems. This limitation arises because BEF/BES studies performed to date have focussed mostly on scavenger community features that ignore functional traits, such as species richness and composition, and abundance of individuals (Mateo-Tomás et al., 2017; Morales-Reyes et al., 2017; Sebastián-González et al., 2016). As a complement to the taxonomic approach, the use of trait-based metrics is a useful framework to better understand the mechanisms driving carrion consumption and the importance of functionally similar species across different geographical areas (Barton et al., 2013; Gravel, Albouy, & Thuiller, 2016; Huijbers et al., 2016; Kane et al., 2017; Kendall, 2013).

Ecological theory provides three main mechanisms that can explain how scavenger community attributes relate to carrion consumption rates at the local scale. First, the “functional identity hypothesis” (Gagic et al., 2015; Grime, 1998; Mokany, Ash, & Roxburgh, 2008) suggests that carrion consumption rates could be enhanced by the presence of key traits within an assemblage. For example, some traits, such as high mobility, large home ranges and information transfer in social vultures, can increase carrion encounter rates (Cortés-Avizanda et al., 2014; Ruxton & Houston, 2004), while others, such as large body sizes (e.g. of large carnivores), can enhance consumption rates through a greater intake capacity (Kane et al., 2017). Second, the “functional diversity hypothesis” (Tilman et al., 1997) posits that carrion consumption rates could increase with trait diversity because niche complementarity increases resource consumption efficiency. Under this hypothesis, scavenger communities showing a more diverse trait representation could increase carrion consumption efficiency through facilitatory interactions and ordered resource partitioning (Hertel & Lehman, 1998; Sebastián-González et al., 2016). For example, corvids, omnivorous and meso-carnivores can help vultures in spotting carcasses, while vultures and large carnivores can open large carcasses, facilitating access for smaller scavengers (König, 1974; Kruuk, 1967). A third hypothesis, the “equivalence hypothesis”, postulates that different individuals and species within a given guild could make equivalent contributions to carrion consumption rates irrespective of their traits (Hubbell, 2005). This hypothesis implies that community aspects, such as scavenger species richness and total abundance or biomass of scavengers, could be the primary determinants of carrion consumption rates.

Using five study areas in Spain and South Africa, characterised by different levels of abundance and richness of vultures, large carnivores and other facultative scavenger species (Mateo-Tomás et al., 2015, 2017), we aimed to identify which scavenger community attributes and ecological mechanisms drive carrion consumption. More specifically, we first explored whether the studied scavenger species could be classified into functionally distinct groups. Second, we quantified different attributes of the scavenger communities that are related to the three hypothesised mechanisms: i) functional identity metrics to reveal functional groups and traits influencing scavenging activity, ii) functional diversity metrics, which describe the range and variety of functional traits within the scavenger community, and iii) metrics representing equivalent contribution of scavenging individuals. Third, to explore which species and community attributes best explain carrion consumption, we correlated the abundance of the main scavenger species, as well as functional identity, diversity and equivalence metrics, with carrion consumption rates. These analyses were conducted at two scales: within each study area and considering all areas together.

## Material and methods

### Study areas

The five study areas were located in temperate (Cordillera Cantábrica, CC) and Mediterranean Spain (Sierra de Cazorla, CZ; Sierra Espuña, ES) and subtropical South Africa (Hluhluwe-iMfolozi Park, HiP; Mkhuze Game Reserve, MK). The study areas represent a gradient of biodiversity, ranging from communities dominated by facultative scavengers to those with a complete scavenger community (i.e. including vultures and large carnivores). While large carnivores are absent from two of the Spanish areas (CZ, ES), CC holds important populations of brown bear (*Ursus arctos*) and wolf (*Canis lupus*). Vultures were common in CZ and CC, but only occasional in ES. The South African areas hold a complete vertebrate scavenger community with large carnivores and vultures, although lions (*Panthera leo*) were absent from MK. Spotted hyaenas (*Crocuta crocuta*) and leopards (*P. pardus*) are common in HiP and MK. *Gyps* species are the most abundant vultures in both Spain (griffon vulture, *G. fulvus*) and South Africa (white-backed vulture, *G. africanus*). Avian and mammalian meso-predators and omnivorous species (e.g. corvids and suids) are common in all areas.

### Scavenger and carcass monitoring

We used motion-triggered remote cameras to monitor 149 carcasses consisting of remains of free-living wild ungulate species obtained from sport hunting, culling and natural mortality in 2006–2013 (see Moleón et al. 2015; Mateo-Tomás et al. 2015 and 2017 for details). The number of carcasses monitored per study area ranged between 11 and 72 (median=19; see Table S1). The large estimated sample coverage for each study area (calculated following Chao et al., 2014) allowed further comparisons of the scavenger communities across the study areas (0.989 - 1.000; see Mateo-Tomás et al., 2017 for additional details on these calculations). Carrion-feeding vertebrates recorded at carcasses were considered the scavenger community (see Mateo-Tomás et al., 2015 for details, 2017).

From the pictures obtained by the camera traps, we calculated *scavenger richness* in each carcass as the number of different species recorded scavenging until complete depletion of the carcass, i.e. when only skin, bones and other hard tissues remained (see Moleón et al., 2015). *Scavenger abundance* was estimated as the highest number of unequivocally different individuals of a given species simultaneously appearing in a picture at each carcass. When possible, we also counted different individuals visiting the same carcass using identifying features like skin patterns, injuries, age, and sexual dimorphism (see Mateo-Tomás et al., 2017). *Scavenger biomass* was estimated as the overall sum of the products between species abundances and their corresponding mean biomass values (as estimated in Mateo-Tomás et al., 2017; Myers et al., 2018; see Supplementary Appendix S1). All of these metrics were calculated for each carcass within each study area.

*Carrion consumption rate* (in kg h^−1^) was calculated at each carcass, as the carcass biomass (in kg) divided by its consumption time (in h). Carcass biomass was measured in the field (using portable scales) and/or by expert assessment and existing references (i.e. when carcasses were too heavy to be weighed *in situ*; see Mateo-Tomás et al., 2017). Consumption time of each carcass was estimated as the time elapsed from carcass placement until its complete depletion (except skin and bones; see above).

### Scavenger community attributes

To explore the main functional attributes of the scavenger community, we characterised the functional traits of the scavenger species recorded at each study area. For this purpose, we compiled a species *x* trait matrix containing information for 13 functional traits that potentially explain differences in consumption rates across species (see Table 1 for traits and their hypothesised relationship with carrion consumption). Trait values were codified based on taxonomic expertise and published information (De Magalhães & Costa, 2009; Kingdon, 2001; Myers et al., 2018; Myhrvold et al., 2015; Skinner & Chimimba, 2005; Wilson & Mittermeier, 2009, 2011). We then used this taxonomic and trait information to quantify three types of community-level metrics across study areas: functional identity, functional diversity and functional equivalence.

**Table 1.**
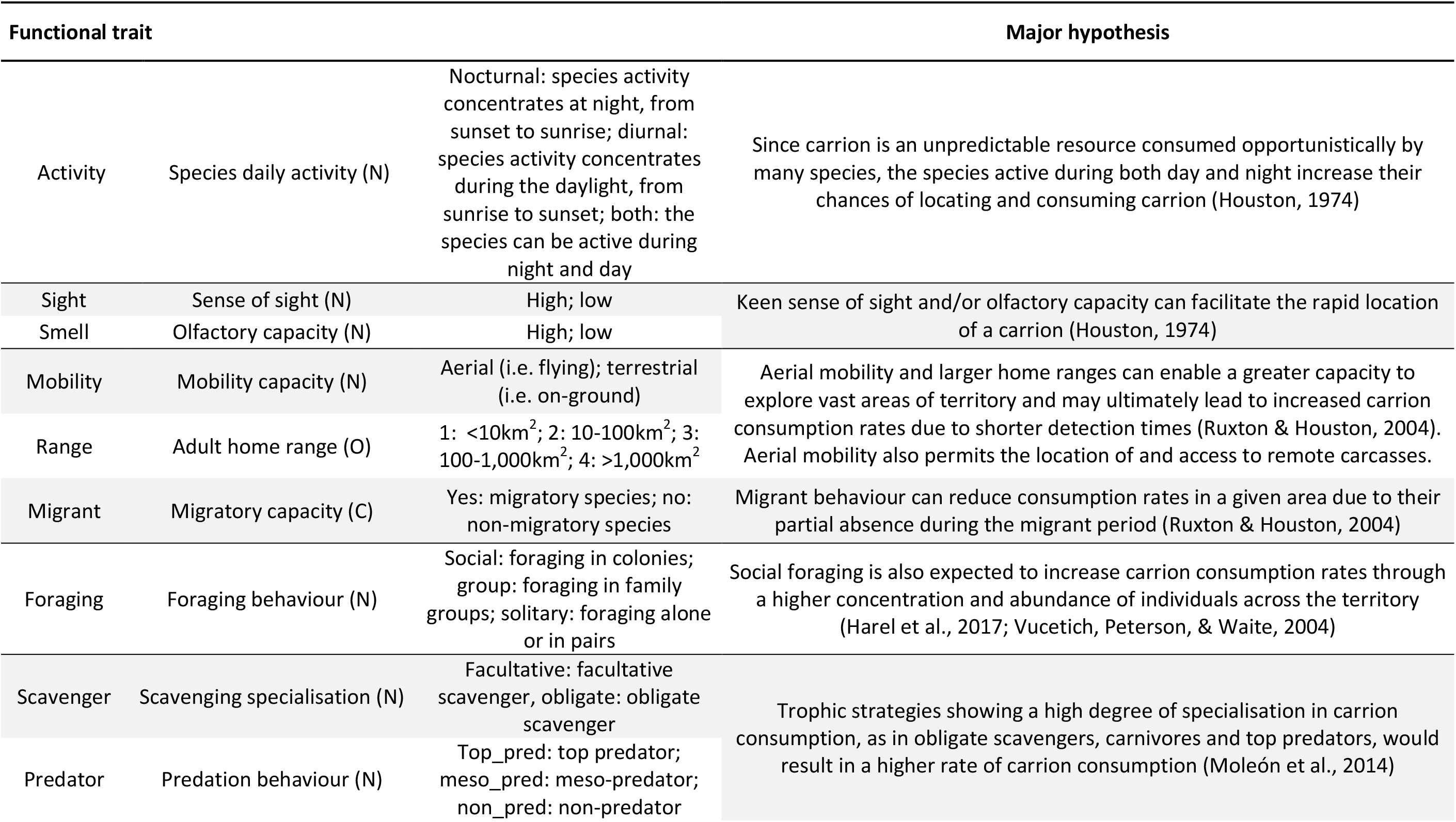
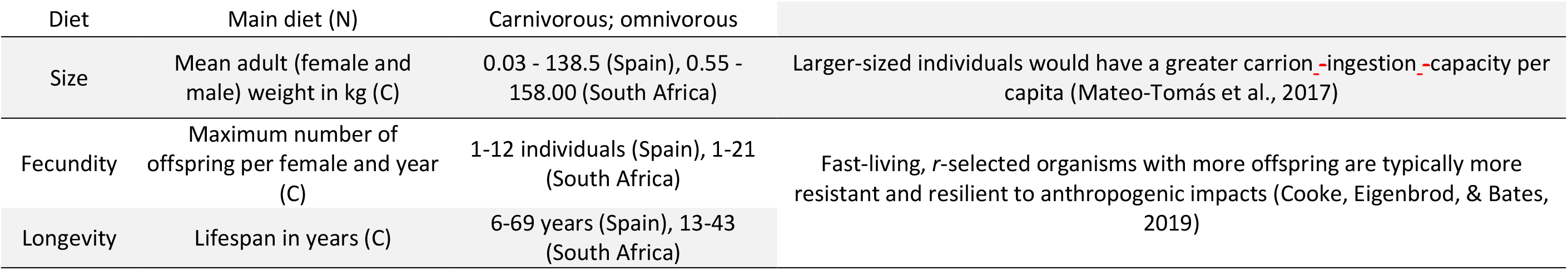
List of functional traits used to characterise scavenger species according to some major hypotheses on carrion consumption. Traits are coded as nominal (N), ordinal (O), or continuous (C) variables, as indicated in parentheses.

First, we estimated two *functional identity* metrics: i) *functional group biomass* and ii) *community mean traits*. On the one hand, we initially computed a Gower dissimilarity matrix based on species traits to obtain functional groups (Gower, 1971), considering all areas together. Then, from this trait dissimilarity matrix, we classified species into a dendrogram (Ward’s clustering method) and pruned it into six groups, as they captured the different scavenger ecological strategies most effectively. Once we identified the main scavenger functional groups, for each study area, we estimated *functional group biomass* as the total biomass of individuals belonging to each functional group, i.e. by summing the products between species abundances at all the monitored carcasses and their corresponding mean biomass values (obtained from literature, Myers et al., 2018). On the other hand, we calculated trait mean values considering the species occurring at each community. These community mean values were weighted by species biomasses. For three categorical traits (foraging behaviour, scavenger specialisation, main diet), we estimated the proportion of species belonging to the trait category that was supposed to be more relevant to driving carrion consumption (proportions of species with social foraging behaviour, obligate scavengers and large carnivores).

Second, we defined two *functional diversity* metrics to describe the variability in functional traits at each study area: *functional dispersion* and *functional richness*. To estimate these metrics, we built a functional space using a Principal Coordinate Analysis (PCoA) from the same Gower trait dissimilarity matrix. We retained the two most explanatory functional axes following Maire et al., (2015) (2D functional space), explaining 79.8% of the trait dissimilarity matrix and showing a good representation of the trait dissimilarity matrix: mean squared deviation of 0.007. For ecological interpretation, we explored the relationship of each functional axis with original traits using Pearson correlation coefficients (quantitative variables) and visual inspection (categorical variables). We calculated functional dispersion (FDis) (Laliberte & Legendre, 2010) as the mean biomass-weighted distance of each species to the community centroid in the 2D functional space. Functional richness (FRic) (Villéger, Mason, & Mouillot, 2008) represents the degree to which each community fills the 2D functional space occupied by all species included in this study (community convex hull divided by all-studied-species convex hull).

Third, we calculated three metrics related to *functional equivalence*: the *richness*, *abundance* and *biomass* of the scavenging species occurring at each community. These metrics are functionally equivalent because they represent indistinguishable species or individual contributions to carrion consumption.

In all cases, species mean biomasses were obtained from the literature (see Supplementary Appendix S1) (Myers et al., 2018).

### Data analyses

We explored which species and community attributes best explained carrion consumption using 1) an exploratory correlation analysis with areas and 2) a model-based hypothesis testing across the five study areas.

First, at the species level, we correlated species abundances and carrion consumption rates within each study area. For the species belonging to each functional group, we averaged correlation values to show if species with similar traits tended to relate similarly to carrion consumption. To avoid spurious correlations, we excluded species with less than five occurrences (i.e. visited carcasses) within a given study area. We also correlated carrion consumption rates with the three types of community-level metrics within each study area: functional group biomass and community mean traits (functional identity); functional dispersion and functional richness (functional diversity); and scavenger richness, abundance and biomass of scavengers (functional equivalence). We used Spearman correlations to avoid problems with data with skewed distributions as a result of subsetting for each area.

Second, we evaluated the support of the functional identity, diversity and equivalence hypotheses in explaining carrion consumption across study areas. For this purpose, we ran a global model on a dataset including all communities and sites (n=149 carcasses), and a group of non-collinear predictors that represent the three competing hypotheses, plus study area as a covariate. To represent the functional identity hypothesis, we selected mean-weighted home range and proportion of large carnivores because they showed the strongest positive relationship to carrion consumption within area correlation and a plausible functional identity mechanism (Table 1). We discarded alternative functional traits or functional group biomasses showing potential predictive power as they all showed a strong collinearity with mean home range or proportion of large carnivores (pairwise Pearson correlation, *r_P_* ≥ |0.70|). As a functional diversity metric, we selected functional dispersion and discarded functional richness, because the latter had a stronger collinearity with functional equivalence predictors (pairwise Pearson correlation, *r_P_* ≥ |0.70|). As a functional equivalence metric, we chose scavenger community abundance because it showed a higher within-area correlation with carrion consumption compared to community biomass and richness. We decided to also include species richness as a functional equivalence metric because it has been traditionally used in BEF studies (Balvanera et al., 2006; Winfree, Fox, Williams, Reilly, & Cariveau, 2015) and enabled us to compare our results to previous works. Selected predictors maintained an acceptable degree of collinearity (pairwise Pearson correlation *r_P_* =-0.03 to *r_P_* =0.69 and variance inflation factor < 3) (Zuur, Ieno, & Elphick, 2009).

Additionally, to explore if community attributes had different effects across study areas, we created five models that included the six predictors (study area and community attributes) plus an interaction term between the study area and one of the five community attributes (one model for each community attribute). We compared each of the models with interactions with a global model without interaction terms using the Akaike’s Information Criterion for small sample sizes (AICc). Only the model including the interaction between species richness and study area was selected over the model without interactions, and therefore this interaction term was included in further analyses. To quantify regression coefficients, statistical support and importance of the selected predictors, we adopted a multi-model inference approach (Burnham, Anderson, & Huyvaert, 2011), using the function *dredge()* from the *MuMIn* R package (Bartoń, 2016). Based on AICc values, this function ranks all the models generated from all the possible combinations of predictors included in our final global model (including the interaction term between richness and study area). All of these models were fitted with a maximum-likelihood method (ML) to allow between-model comparisons. We retained all the ranked models accounting for a 95% cumulative Akaike weight. To visualise the overall response of these models, we averaged their predictions weighing by each model’s Akaike weight. To approximate the contribution of each predictor to total variance in the model ranked first (minimum AICc), we partitioned their explained variance by calculating the unique contribution to explained variance of each predictor. For averaged predictions and variance partition, models were re-fitted by maximising the restricted log-likelihood (REML). All models were performed using General Least Squares (GLS) with a variance structure to account for between-study area variance heterogeneity (heteroscedasticity) and were validated by visually checking their residuals for normality and homoscedasticity. Before analyses, to reduce variable distribution skewness and improve linearity, we applied a square-root-transformation to functional dispersion, a log-transformation to carrion consumption rate, abundance, biomass, and all functional group biomasses, and a logit-transformation to all mean-weighted proportions of trait categories. All the quantitative predictors were also standardised to mean=0 and SD=1 to facilitate model coefficient comparisons. The code and functions used to run all of these analyses are available in Supplementary Appendix S1, and were conducted using R version 3.5.2 (R Development Core Team, 2019).

## Results

### Scavengers’ functional groups

Our functional classification revealed six clearly distinct ecological strategies (Fig. 1a). Three functional groups (corvids, vultures and other raptors) included avian species with aerial mobility, medium to high longevity (13-48 years), low fecundity (1-9 offspring per year), low olfactory capacity and strong sense of sight. The other three groups (omnivores, meso-carnivores and large carnivores) represented terrestrial mammals with facultative scavenging, low to medium longevity (6-45 years), low sense of sight and high fecundity (1.5-21 offspring per year) and olfactory capacity. Corvids and other raptors were facultative scavengers. Corvids accounted for small-sized omnivorous birds (0.2-2.0 kg) with small home ranges (<10 km^2^), while other raptors included small- to medium-sized avian predators (eagles and kites; 0.6-4.0 kg) with small to intermediate home ranges (<10 to 10-100 km^2^). The vultures’ group was made up of obligate scavengers of intermediate body-size (2.0-9.8 kg), large to very large home ranges (100-1000 to >1000 km^2^) and variable social behaviour. The omnivorous group included large-sized (52.98-71.00 kg) suids with low to intermediate home ranges (<10 to 10-100 km^2^) and a small (0.02 kg) rodent with a small home range (<10 km^2^). Meso-carnivores included mammalian meso-predators, carnivores of intermediate size (1.7-15.0 kg) and low to intermediate home range (<10 to 10-100 km^2^), while the large carnivore group comprised top predators, large-sized organisms (26.0-158.0 kg), all with large home ranges (100-1000 km^2^) but with both carnivorous (felids, wolf) or omnivorous diets (bear). These groups were distributed across two dimensions of trait variation (functional space) that, in combination, explained 79.8% of the original trait variation (Axis 1: 63.8%, Axis 2: 16.0%; Fig. 1b).

**Fig. 1.**
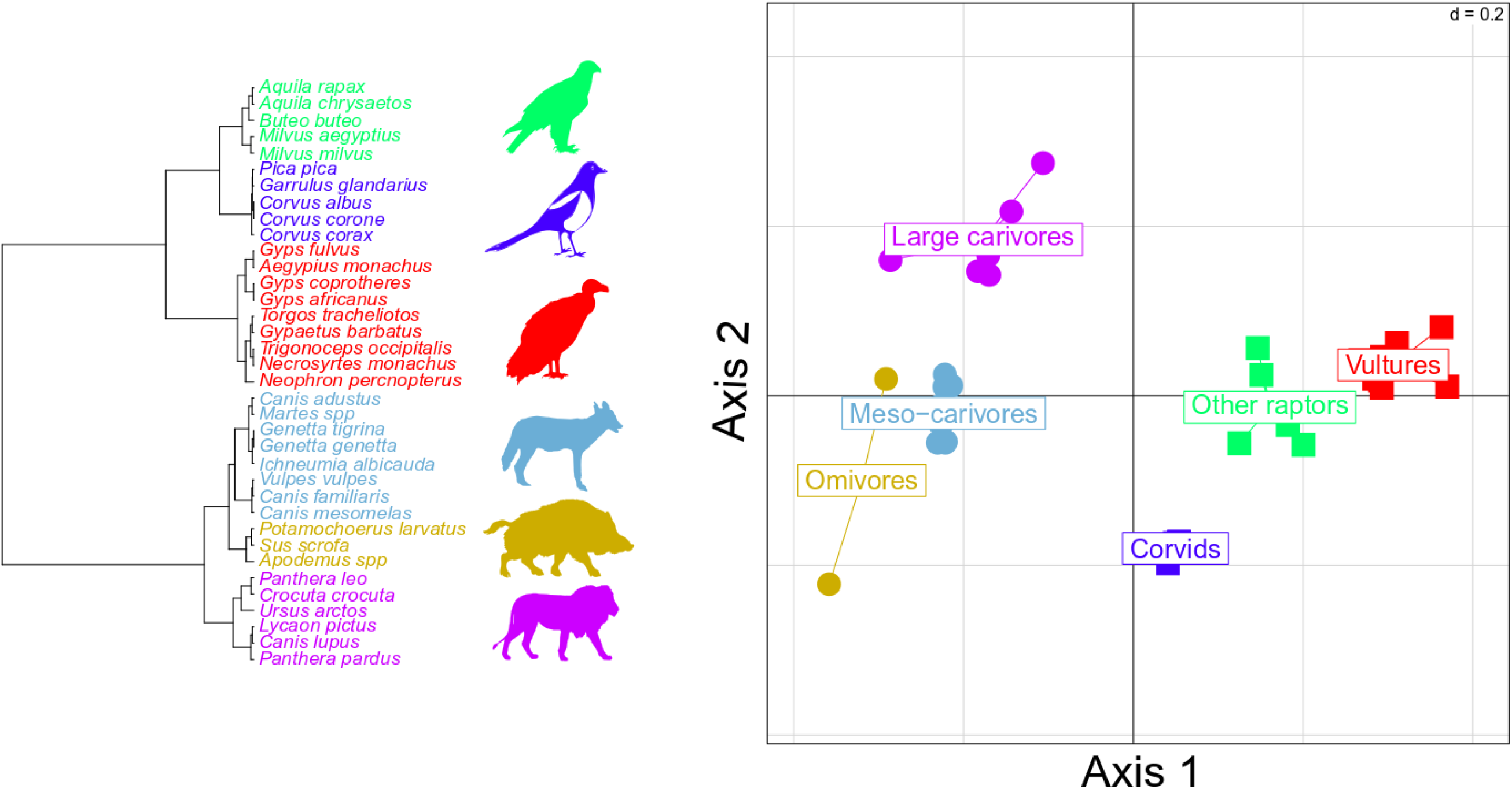
Cluster showing the functional classification of the scavengers (a) and two first axes of the functional space showing the functional similarity across scavenger species (b). The functional groups are represented by different colours: corvids (green), other raptors (dark blue), vultures (red), omnivores (golden), meso-carnivores (light blue) and large carnivores (purple). Axis 1’s positive values were linked to avian organisms with aerial mobility, an omnivorous diet, diurnal activity, good sight ability and high longevity (*r*=0.54), while negative values were associated with terrestrial mammal organisms with a carnivorous diet, nocturnal or both nocturnal and diurnal activity, high olfactory capacity and high fecundity (*r*=-0.69). Positive values of axis 2 were related to carnivores of large size (*r*=0.82), large home ranges (*r*=0.74) and high longevity (*r*=0.52).

### Within-area drivers of carrion consumption

The abundances of the functional groups of vultures and large carnivores generally showed the highest positive correlations with carrion consumption rates (Fig. 2a), while the abundances of species featuring different strategies, such as meso-carnivores and omnivorous functional groups, showed a more frequent number of negative correlations (Fig 2a, Table S3). The abundance of griffon vulture (*ρ*=0.50 to 0.63; CC and CZ) was highly and positively correlated with carrion consumption rates (Fig. S3, Table S2). A second group of species showed weaker positive correlations between abundance and carrion consumption rates, including the lion (*ρ*=0.20) and the golden eagle (*Aquila chrysaetos, ρ*=0.12 to 0.28). A third group of species exhibited a study area-dependent effect, resulting in highly variable correlation values between abundance and carrion consumption rates. For example, red fox abundance showed a positive association to carrion consumption rate only in ES (*ρ*=0.41), but negative correlations in the other Spanish areas (*ρ*=-0.31 to −0.25). A fourth group of organisms, made up of the omnivore and meso-carnivore functional groups, showed a consistent negative correlation between abundance and carrion consumption rates, and included the wild dog (*C. familiaris, ρ*=-0.51 to −0.28), bush pig (*Potamochoerus larvatus, ρ*=-0.38), Cape genet (*Genetta tigrina, ρ*=-0.47) and red kite (*Milvus milvus, ρ*=-0.24 to −0.19).

**Fig. 2.**
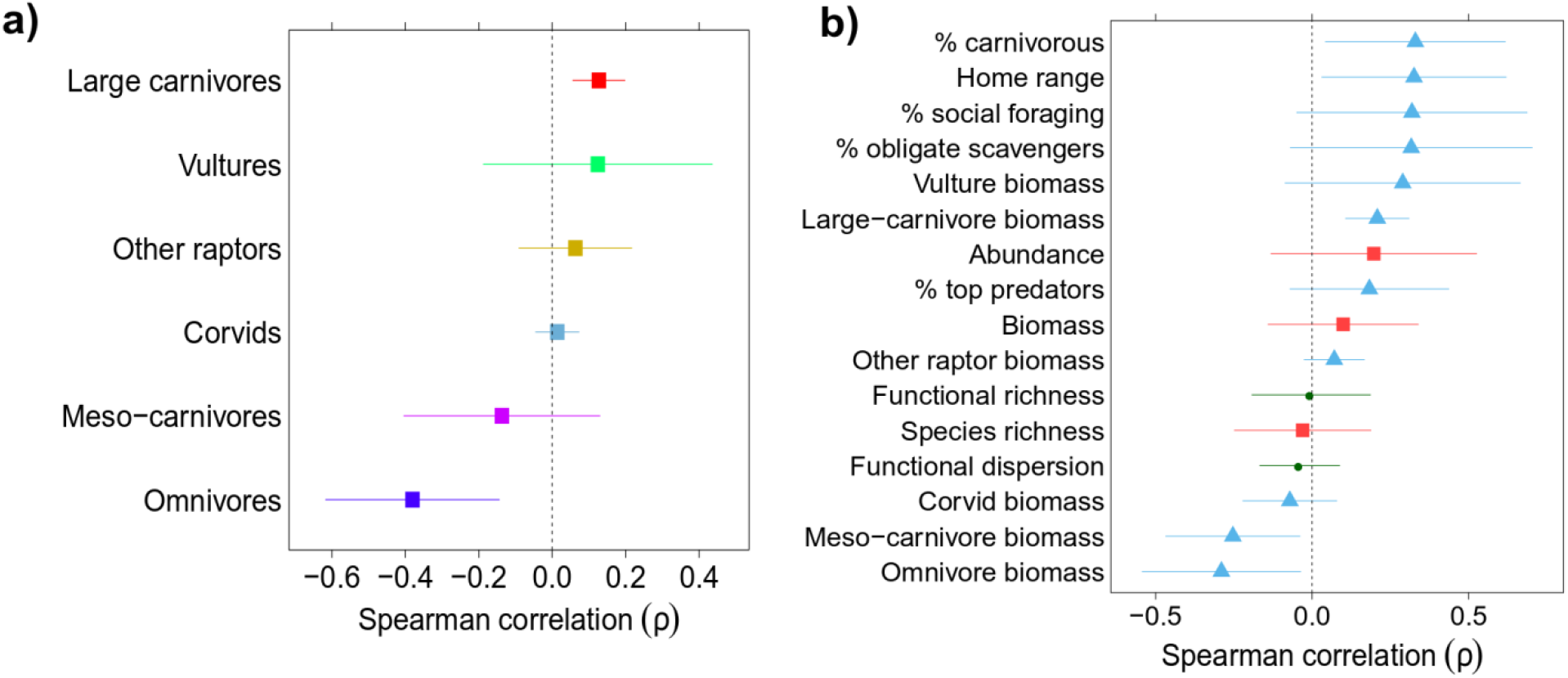
Mean (± 95% CI) of within-area Spearman correlations between carrion consumption rates and species abundances at each carcass, classified into functional groups (a), and community-level attributes (b). Colours in (a) as in Fig.1, while colours in (b) represent the three types of community-level metrics: functional identity (blue triangles), functional diversity (green circles) and functional equivalence (red squares). See Tables S2 and S3 for more details.

The proportion of species with a carnivorous diet (Spearman correlation, *ρ*=-0.14 to *ρ*=0.71) and mean home range (*ρ*=-0.16 to *ρ*=0.67) within the community showed the highest positive associations to carrion consumption in most study areas, followed by the proportion of social foraging species (*ρ*=-0.17 to *ρ*=0.69), the proportion of obligate scavengers (*ρ*=-0.22 to *ρ*=0.69) and the biomass of vultures (*ρ*=-0.24 to *ρ*=0.64) and large carnivores (*ρ*=0.12 to *ρ*=0.30) (Fig. 2b, Table S3). Mean home range and the proportion of species with a carnivorous diet showed the highest positive values for CZ, CC and MK. Abundance and biomass showed both positive and negative relationships to carrion consumption rates (*ρ*=-0.34 to *ρ*=0.60). Noticeably, abundance showed a high positive correlation in CZ (*ρ*=0.60). Functional richness, functional dispersion and species richness showed overall weak correlations to carrion consumption rate in all study areas (*ρ*=-0.31 to *ρ*=0.29). The biomass of omnivore (*ρ*=-0.60 to *ρ*=0.02) and meso-carnivore (*ρ*=-0.45 to *ρ*=0.03) groups exhibited the most negative mean correlation to carrion consumption rate. Some community attributes varied in importance according to the study area. For example, traits related to obligate scavengers, such as the proportion of obligate scavengers or social foraging individuals, showed much higher correlation coefficients in CZ and CC (*ρ*=0.59 to *ρ*=0.61) than in MK and HiP (*ρ*=-0.17 to *ρ*=0.28).

### Between-area drivers of carrion consumption

We found 21 models within the 95% cumulative Akaike weight range (Table 2 and Table S4). Home range was included in all of these models and showed a positive correlation to carrion consumption rate (coefficient range: 0.68 to 0.80) when the five study areas were considered together (Table 2, Fig. 3a). Species richness was included in 14 out of the 21 best models and had an overall negative effect on carrion consumption rates (coefficient range: −0.31 to −0.08; Fig. 3b). The proportion of large carnivores was included in seven models, having a weak negative effect in five of them (coefficient range: −0.06 to 0.00). Functional dispersion (coefficient range: −0.02 to 0.18) and community abundance (coefficient range: −0.07 to 0.11) were included in ten and nine models, respectively, and showed positive slopes in most (seven and six models, respectively). Carcass weight was included in all the models and had a positive effect on carrion consumption (coefficient range: 0.41 to 0.60; Table 2). The best model (i.e. the one with the lowest AICc value) included mean home range, species richness and carcass weight as predictors (Table 2). This model accounted for 48.1% of the variation of the carrion consumption rate, with mean home range (17.9%) showing a much higher explanatory capacity than species richness (0.1%). Carcass weight explained 6.4% of the variance, and the term to reduce variance heterogeneity between study areas accounted for 23.7% of variance.

**Table 2.**
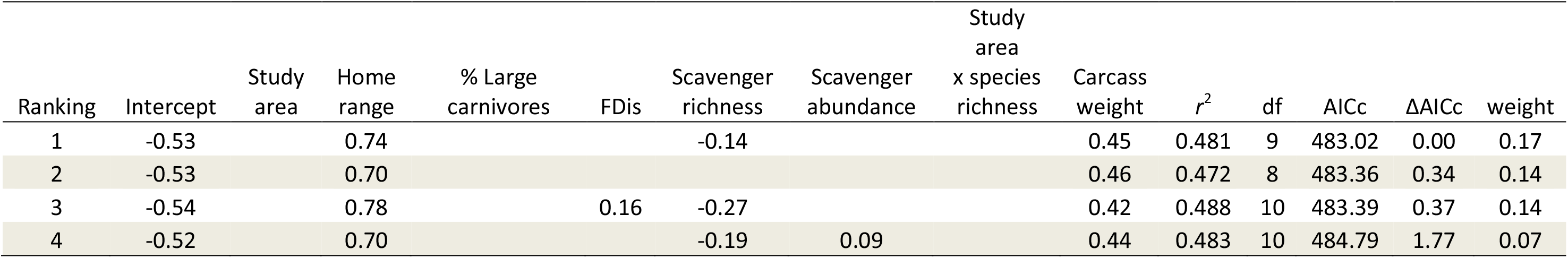
Top four models relating carrion consumption rate to study area and selected scavenger community attributes: home range, proportion of large carnivores (% large carnivores), functional dispersion (FDis), scavenger species richness and abundance. Carcass weight was also included as a predictor. Regression coefficients, goodness-of-fit (*r*^2^), degrees of freedom (df), AICc and ΔAICc values, and model weight are also shown. +: area / interaction terms were included in the model. Blank spaces indicate that the predictor was not included in the model; see full results in Table S4.

**Fig 3.**
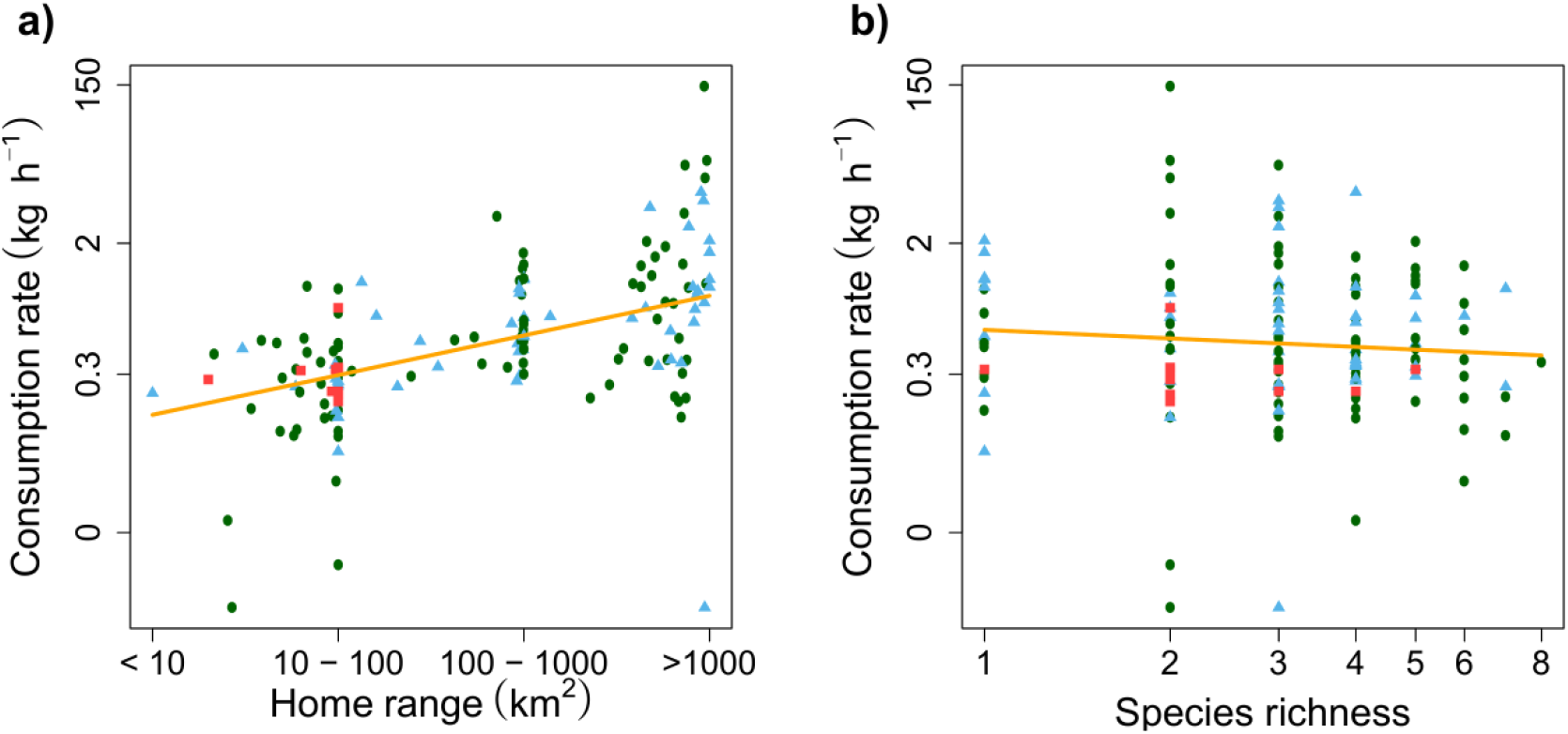
Variation in consumption rate in relation to mean home range (a) and species richness (b) across study areas. Fitted values were calculated through averaging model predictions weighted by each model’s Akaike weight. Carcasses (points) belonging to different study areas are represented by different symbols depending on their scavenger composition. F (red squares): only facultative scavengers (ES); FO (green circles): facultative + obligate (CZ and MK); FOP (blue triangles): facultative + obligate + large carnivores (CC and HiP).

## Discussion

In accordance with the functional identity hypothesis, our results revealed that scavenging vertebrates with larger home ranges support higher rates of carrion consumption. In contrast, the functional diversity and functional equivalence hypotheses had a generally poor explanatory capacity. Thus, the identity hypothesis, which assumes the dominant role of key traits in driving ecosystem functioning (Gagic et al., 2015; Grime, 1998; Mokany et al., 2008), emerges as a better framework for explaining carrion consumption rates than niche complementarity (i.e. functional diversity; Tilman et al. 1997) or equivalence (i.e. species richness or abundance; Hubbell, 2005). Interestingly, the relationship between carrion consumption rate and scavenger home range was consistent to varying defaunation degrees (i.e. from less to more diverse scavenger communities), irrespective of the species and functional groups present at each study area and the carcass size

Large home ranges, such as those seen in vultures (e.g. >1,000 km^2^), can enhance carrion consumption through an increased capacity to detect carcasses (i.e. reducing detection time) over vast territories as opposed to the smaller foraging areas of facultative scavengers (Cortés-Avizanda, Jovani, Donázar, & Grimm, 2012; Kane et al., 2017; Ruxton & Houston, 2004). In the case of vultures, aerial mobility through soaring flight also confers rapid access to carcasses with a low energy expenditure in comparison to terrestrial mammals and non-soaring birds (Ruxton & Houston, 2004). In addition, species with a large home range generally have other attributes that confer increased scavenging efficiency and competitive abilities, such as larger body sizes (Devault et al., 2003; Moleón et al., 2014; Ruxton & Houston, 2004). Another common feature in scavenging species with large home ranges, especially *Gyps* vultures and large carnivores such as lions, hyaenas and wolves, is an enhanced social behaviour, which allows the efficient communication of information across large spans of territory (Harel, Spiegel, Getz, & Nathan, 2017; Kruuk, 1972) and an increased intra-specific recruitment at carcasses (Cortés-Avizanda et al., 2014). Species with enhanced social behaviour can also monopolise carcasses more easily and avoid interspecific competition (Cortés-Avizanda et al., 2012), thereby increasing scavenging efficiency. Indeed, our variable depicting social foraging was associated with rapid consumption rates.

Most of the previous BEF/BES research has generally found a positive effect of species richness on ecosystem functioning (Balvanera et al., 2006; Duncan et al., 2015). However, in our study, we found that species richness was a poor predictor of carrion consumption. In a recent study, Mateo-Tomás et al. (2017) showed that species richness increased carrion consumption only in areas where functionally dominant species, such as some obligate scavengers (i.e. *Gyps* vultures) and large carnivores (i.e. lions and hyenas), were rare. The poor predictive capacity of species richness in our study suggests that functional equivalence (i.e. niche equivalence, mass effect) is a less suitable and generalisable framework to explain carrion consumption than functional identity metrics, such as those describing the abundance of species with key traits. Some previous studies also found a higher explanatory capacity of functional identity metrics (Gagic et al., 2015; Mokany et al., 2008; Winfree et al., 2015). Moreover, our data suggest that efficient carrion consumption was not enhanced by complementary and diverse traits in scavenger communities, as indicated by the low explanatory capacity of functional diversity metrics such as functional richness or dispersion. This contrasts with evidence showing bidirectional facilitatory processes between small-sized facultative scavengers and obligate scavengers (e.g. functional groups such as corvids, omnivores and meso-carnivores can help vultures spot carcasses) and large carnivores (that can open large carcasses and facilitate access to smaller scavengers) (König, 1974; Kruuk, 1967). This lack of complementarity could result from the spatio-temporal partitioning of the carcasses by different species (Blázquez, Sánchez-Zapata, Botella, Carrete, & Eguía, 2009) and may perhaps be due to the reduced carrion intake capacity of most generalists (but see Mateo-Tomás et al., 2017). However, the ecological mechanisms linking species and trait richness and carrion consumption may change at regional scales (Mateo-Tomás et al., 2017). For example, consumption rates may be enhanced in highly heterogeneous landscapes if communities show complementary and diverse traits adapted to the detection of carcasses under different habitats.

Large carnivores are key scavengers in the South African areas, as reflected by their high consumption rates and total carrion consumptions (Mateo-Tomás et al., 2017). Their greater importance in South Africa could be a result of their greater abundance compared to areas in Spain, and their competitive superior abilities when directly confronting African vultures (Kendall, 2013; Kendall, Virani, Kirui, Thomsett, & Githiru, 2012). In addition, one of the two large carnivore species present in Spain, the brown bear, is a solitary species that feeds on a very wide variety of food resources, including important quantities of plant material (Naves, Fernández-Gil, Rodríguez, & Delibes, 2006). This particular behaviour may have contributed to explaining the high efficiency in carrion consumption shown by vultures in Spain, even in an area with large carnivores. In areas without large carnivores and vultures, other scavengers with relatively large home ranges, such as eagles, foxes and suids, became the most efficient scavengers (Mateo-Tomás et al., 2017).

Despite the general assumption that large-bodied organisms play a key role in sustaining ecosystem functions (Doughty et al., 2016; Galetti et al., 2018), few studies have identified which particular attributes and mechanisms explain the functional role of terrestrial vertebrate communities (Duncan et al., 2015; Sobral et al., 2017). As a main advantage, trait-based metrics offer a complementary argument for biodiversity conservation and the maintenance of ecosystem functions, complementing traditional approaches based on species richness or abundance. For example, our case study suggests that policies focussed on scavengers with large home ranges may be an effective target for management programs aimed at enhancing the scavenging ecological function. However, it is necessary to investigate to what degree carrion consumption can be sustained at moderate or high levels without compromising other key functions, such as pest control or seed dispersal (i.e. multifunctionality), due to unavoidable trade-offs, as suggested in previous research (e.g. Felipe-Lucia et al., 2018). Further research should explore to what degree organisms with large home ranges, and thus the functional identity hypothesis framework, are critical to sustaining carrion consumption in a more general context, i.e. including different carcass sizes, seasons and ecosystem functions, as well as potential interactions with invertebrate scavengers.

In conclusion, we showed that the functional identity framework explained carrion removal patterns across different areas better than alternative approaches considering species complementarity and equivalence. In particular, species home range size seems to be a key characteristic in predicting carrion removal rates. Our results also reinforce the idea of the pivotal role of mobile link organisms in ecosystem functioning (Lundberg & Moberg, 2003), which has already been highlighted for vultures in scavenger assemblages (Mateo-Tomás, Olea, Selva, & Sánchez-Zapata, 2019), and as providers of ecosystem services at the landscape level (Kremen, Williams, Bugg, Fay, & Thorp, 2004). Most of these links have been explored for specific taxonomic guilds and functions (i.e. birds and dispersal, bees and pollination). In contrast, our guild of vertebrate scavengers includes a heterogeneous taxonomic composition with contrasting traits. Among those traits, the home range of vertebrate scavengers is particularly variable with species moving from hundreds of metres to thousands of km. Thus scavenger guilds may provide a paradigmatic framework to explore ecosystem processes in relation to movement ecology in a changing world (Donázar et al., 2016).

## Supporting information

File including supplementary tables and figures

Files including R functions and code to reproduce some of our statistical analyses

## Acknowledgements

CG-C and ESG were supported by “Juan de la Cierva” research contracts (MINECO, FJCI-2015-25785 & IJCI-2015-24947, respectively), and MM by a postdoctoral fellowship from the Spanish Ministry of Education (Plan Nacional de I+D+i 2008-2011), by the Severo Ochoa Program for Centres of Excellence in R+D+I (SEV-2012-0262) and by a research contract Ramón y Cajal from the MINECO (RYC-2015-19231). CG-C was also supported by the European Regional Development Fund (COMPETE2020 and PT2020) and the Portuguese Foundation for Science and Technology (FCT), through the Centre of Molecular and Environmental Biology (CBMA) strategic program UID/BIA/04050/2019 (POCI-01-0145-FEDER-007569) and the STREAMECO project (Biodiversity and ecosystem functioning under climate change: from the gene to the stream, PTDC/CTA-AMB/31245/2017). ESG was also supported by Generalitat Valenciana (SEJI/2018/024). This study was partly funded by the Spanish Ministry of Economy, Industry and Competitiveness and EU ERDF funds through the projects CGL2015-66966-C2-1-2-R and CGL2017-89905-R.

## References

Balvanera, P., Pfisterer, A. B., Buchmann, N., He, J. S., Nakashizuka, T., Raffaelli, D., & Schmid, B. (2006). Quantifying the evidence for biodiversity effects on ecosystem functioning and services. Ecology Letters, 9(10), 1146–1156. doi:10.1111/j.1461-0248.2006.00963.x

Bartoń, K. (2016). MuMIn: Multi-model inference. R package version 1.15.6. Version, 1, 18.

Barton, P. S., Cunningham, S. A., Macdonald, B. C. T., McIntyre, S., Lindenmayer, D. B., & Manning, A. D. (2013). Species traits predict assemblage dynamics at ephemeral resource patches created by carrion. PLoS ONE, 8(1). doi:10.1371/journal.pone.0053961

Beasley, J. C., Olson, Z. H., Selva, N., & DeVault, T. L. (2019). Ecological functions of vertebrate scavenging. In P. P. Olea, P. Mateo-Tomás, & J. A. Sánchez Zapata (Eds.), Carrion ecology and management (pp. 125–157). Cham (Switzerland): Springer International Publishing.

Blázquez, M., Sánchez-Zapata, J. A., Botella, F., Carrete, M., & Eguía, S. (2009). Spatio-temporal segregation of facultative avian scavengers at ungulate carcasses. Acta Oecologica, 35(5), 645–650. doi:10.1016/j.actao.2009.06.002

Burnham, K. P., Anderson, D. R., & Huyvaert, K. P. (2011). AIC model selection and multimodel inference in behavioral ecology: Some background, observations, and comparisons. Behavioral Ecology and Sociobiology, 65(1), 23–35. doi:10.1007/s00265-010-1029-6

Chao, A., Gotelli, N. J., Hsieh, T. C., Sander, E. L., Ma, K. H., Colwell, R. K., & Ellison, A. M. (2014). Rarefaction and extrapolation with Hill numbers: A framework for sampling and estimation in species diversity studies. Ecological Monographs, 84(1), 45–67. doi:10.1890/13-0133.1

Cooke, R. S. C., Eigenbrod, F., & Bates, A. E. (2019). Projected losses of global mammal and bird ecological strategies. Nature Communications, 10(1), 2279. doi:10.1038/s41467-019-10284-z

Cortés-Avizanda, A., Jovani, R., Donázar, J. A., & Grimm, V. (2012). Resource unpredictability promotes species diversity and coexistence in an avian scavenger guild: a field experiment. Ecology, 93(12), 2570–2579.

Cortés-Avizanda, A., Jovani, R., Donázar, J. A., & Grimm, V. (2014). Bird sky networks: How do avian scavengers use social information to find carrion? Ecology, 95(7), 1799–1808. doi:10.1890/13-0574.1

De Magalhães, J. P., & Costa, J. (2009). A database of vertebrate longevity records and their relation to other life-history traits. Journal of Evolutionary Biology, 22(8), 1770–1774. doi:10.1111/j.1420-9101.2009.01783.x

Devault, T. L., Rhodes, O. E., Shivik, J. A., & Ewjtt, X. J. (2003). Scavenging by vertebrates: behavioral, ecological, and evolutionary perspectives on an important energy transfer pathway in terrestrial ecosystems. Oikos, 102(2), 225–234.

Donázar, J. A., Cortés-Avizanda, A., Fargallo, J. A., Margalida, A., Moleón, M., Morales-Reyes, Z., … Serrano, D. (2016). Roles of raptors in a changing world: from flagships to providers of key ecosystem services. Ardeola, 63, 181–234. doi:10.13157/arla.63.1.2016.rp8

Doughty, C. E., Roman, J., Faurby, S., Wolf, A., Haque, A., Bakker, E. S., … Svenning, J.-C. (2016). Global nutrient transport in a world of giants. Proceedings of the National Academy of Sciences, 113(4), 868–873.

Duncan, C., Thompson, J. R., & Pettorelli, N. (2015). The quest for a mechanistic understanding of biodiversity–ecosystem services relationships. Proceedings of the Royal Society B: Biological Sciences, 282(1817), 20151348. doi:10.1098/rspb.2015.1348

Felipe-Lucia, M. R., Soliveres, S., Penone, C., Manning, P., van der Plas, F., Boch, S., … Allan, E. (2018). Multiple forest attributes underpin the supply of multiple ecosystem services. Nature Communications, 9, 4839. doi:10.1038/s41467-018-07082-4

Gagic, V., Bartomeus, I., Jonsson, T., Taylor, A., Winqvist, C., Fischer, C., … Bommarco, R. (2015). Functional identity and diversity of animals predict ecosystem functioning better than species-based indices. Proceedings of the Royal Society B: Biological Sciences, 282, 20142620. doi:10.1098/rspb.2014.2620

Galetti, M., Moleón, M., Jordano, P., Pires, M. M., Guimarães, P. R., Pape, T., … Svenning, J. C. (2018). Ecological and evolutionary legacy of megafauna extinctions. Biological Reviews, 93(2), 845–862. doi:10.1111/brv.12374

Gower, J. C. (1971). A general coefficient of similarity and some of its properties. Biometrics, 27(4), 857–871. doi:10.2307/2528823

Gravel, D., Albouy, C., & Thuiller, W. (2016). The meaning of functional trait composition of food webs for ecosystem functioning. Philosophical Transactions of the Royal Society B: Biological Sciences, 371(1694). doi:10.1098/rstb.2015.0268

Grime, J. P. (1998). Benefits of plant diversity to ecosystems: Immediate, filter and founder effects. Journal of Ecology, 86(6), 902–910. doi:10.1046/j.1365-2745.1998.00306.x

Harel, R., Spiegel, O., Getz, W. M., & Nathan, R. (2017). Social foraging and individual consistency in following behaviour: Testing the information centre hypothesis in free-ranging vultures. Proceedings of the Royal Society B: Biological Sciences, 284(1852). doi:10.1098/rspb.2016.2654

Hertel, F., & Lehman, N. (1998). A randomized nearest-neighbor approach for assessment of character displacement: The vulture guild as a model. Journal of Theoretical Biology, 190, 51–61. doi:10.1006/jtbi.1997.0531

Hooper, D. U., Adair, E. C., Cardinale, B. J., Byrnes, J. E. K., Hungate, B. A., Matulich, K. L., … O’Connor, M. I. (2012). A global synthesis reveals biodiversity loss as a major driver of ecosystem change. Nature, 486, 105–108. doi:10.1038/nature11118

Houston, D. C. (1974). Food searching in griffon vultures. African Journal of Ecology, 12(1), 63–77. doi:10.1111/j.1365-2028.1974.tb00107.x

Hubbell, S. P. (2005). Neutral theory in community ecology and the hypothesis of functional equivalence. Functional Ecology, 19(1), 166–172. doi:10.1111/j.0269-8463.2005.00965.x

Huijbers, C. M., Schlacher, T. A., McVeigh, R. R., Schoeman, D. S., Olds, A. D., Brown, M. B., … Connolly, R. M. (2016). Functional replacement across species pools of vertebrate scavengers separated at a continental scale maintains an ecosystem function. Functional Ecology, 30(6), 998–1005. doi:10.1111/1365-2435.12577

Inger, R., Per, E., Cox, D. T. C., & Gaston, K. J. (2016). Key role in ecosystem functioning of scavengers reliant on a single common species. Scientific Reports, 6, 2–6. doi:10.1038/srep29641

Kane, A., Healy, K., Guillerme, T., Ruxton, G. D., & Jackson, A. L. (2017). A recipe for scavenging in vertebrates – the natural history of a behaviour. Ecography, 40(2), 324–334. doi:10.1111/ecog.02817

Kendall, C. J. (2013). Alternative strategies in avian scavengers: How subordinate species foil the despotic distribution. Behavioral Ecology and Sociobiology, 67(3), 383–393. doi:10.1007/s00265-012-1458-5

Kendall, C. J., Virani, M. Z., Kirui, P., Thomsett, S., & Githiru, M. (2012). Mechanisms of Coexistence in Vultures: Understanding the Patterns of Vulture Abundance at Carcasses in Masai Mara National Reserve, Kenya. The Condor, 114(3), 523–531. doi:10.1525/cond.2012.100196

Kingdon, J. (2001). A database of vertebrate longevity records and their relation to other life-history traits.

König, C. (1974). Zum Verhalten spanischer Geier an Kadavern. Journal of Ornithology, 115(3), 289–320. doi:10.1007/BF01644326

Kremen, C., Williams, N. M., Bugg, R. L., Fay, J. P., & Thorp, R. W. (2004). The area requirements of an ecosystem service: Crop pollination by native bee communities in California. Ecology Letters, 7, 1109–1119. doi:10.1111/j.1461-0248.2004.00662.x

Kruuk, H. (1967). Commpetition for food between vultures in East Africa. Ardea, 55, 171–193.

Kruuk, H. (1972). The Spotted Hyaena. A Study of Predation and Social Behavior. Chicago: University of Chicago Press.

Laliberte, E., & Legendre, P. (2010). A distance-based framework for measuring functional diversity from multiple traits. Ecology, 91(1), 299–305. doi:10.1890/08-2244.1

Lundberg, J., & Moberg, F. (2003). Mobile link organisms and ecosystem functioning: Implications for ecosystem resilience and management. Ecosystems, 6, 87–98. doi:10.1007/s10021-002-0150-4

Maire, E., Grenouillet, G., Brosse, S., & Villéger, S. (2015). How many dimensions are needed to accurately assess functional diversity? A pragmatic approach for assessing the quality of functional spaces. Global Ecology and Biogeography, 24(6), 728–740. doi:10.1111/geb.12299

Markandya, A., Taylor, T., Longo, A., Murty, M. N., Murty, S., & Dhavala, K. (2008). Counting the cost of vulture decline-An appraisal of the human health and other benefits of vultures in India. Ecological Economics, 67(2), 194–204. doi:10.1016/j.ecolecon.2008.04.020

Mateo-Tomás, P., Olea, P. P., Moleón, M., Selva, N., & Sánchez-Zapata, J. A. (2017). Both rare and common species support ecosystem services in scavenger communities. Global Ecology and Biogeography, 26(12), 1459–1470. doi:10.1111/geb.12673

Mateo-Tomás, P., Olea, P. P., Moleón, M., Vicente, J., Botella, F., Selva, N., … Sánchez-Zapata, J. A. (2015). From regional to global patterns in vertebrate scavenger communities subsidized by big game hunting. Diversity and Distributions, 21(8), 913–924. doi:10.1111/ddi.12330

Mateo-Tomás, P., Olea, P. P., Selva, N., & Sánchez-Zapata, J. A. (2019). Species and individual replacements contribute more than nestedness to shape vertebrate scavenger metacommunities. Ecography, 42(2), 365–375. doi:10.1111/ecog.03854

Mokany, K., Ash, J., & Roxburgh, S. (2008). Functional identity is more important than diversity in influencing ecosystem processes in a temperate native grassland. Journal of Ecology, 96(5), 884–893. doi:10.1111/j.1365-2745.2008.01395.x

Moleón, M., Sánchez-Zapata, J. A., Sebastián-González, E., & Owen-Smith, N. (2015). Carcass size shapes the structure and functioning of an African scavenging assemblage. Oikos, 124(10), 1391–1403. doi:10.1111/oik.02222

Moleón, M., Sánchez-Zapata, J. A., Selva, N., Donázar, J. A., & Owen-Smith, N. (2014). Inter-specific interactions linking predation and scavenging in terrestrial vertebrate assemblages. Biological Reviews, 89(4), 1042–1054. doi:10.1111/brv.12097

Morales-Reyes, Z., Sánchez-Zapata, J. A., Sebastián-González, E., Botella, F., Carrete, M., & Moleón, M. (2017). Scavenging efficiency and red fox abundance in Mediterranean mountains with and without vultures. Acta Oecologica, 79, 81–88. doi:10.1016/j.actao.2016.12.012

Myers, P., Espinosa, R., Parr, C. S., Jones, T., Hammond, G. S., & Dewey, T. A. (2018). The Animal Diversity Web.

Myhrvold, N. P., Baldridge, E., Chan, B., Sivam, D., Freeman, D. L., & Ernest, S. K. M. (2015). An amniote life-history database to perform comparative analyses with birds, mammals, and reptiles. Ecology, 95(11), 3109–3109. doi:10.1890/15-0846r.1

Naves, J., Fernández-Gil, A., Rodríguez, C., & Delibes, M. (2006). Brown bear food habits at the border of its range: a long-term study. Journal of Mammalogy, 87(5), 899–908. doi:10.1644/05-mamm-a-318r2.1

Pereira, L. M., Owen-Smith, N., & Moleón, M. (2014). Facultative predation and scavenging by mammalian carnivores: Seasonal, regional and intra-guild comparisons. Mammal Review, 44(1), 44–55. doi:10.1111/mam.12005

R Development Core Team. (2019). R Development Core Team. R: A language and environment for statistical computing. Vienna, Austria: R Foundation for Statistical Computing. Retrieved from https://www.r-project.org/

Ruxton, G. D., & Houston, D. C. (2004). Obligate vertebrate scavengers must be large soaring fliers. Journal of Theoretical Biology, 228(3), 431–436. doi:10.1016/j.jtbi.2004.02.005

Sebastián-González, E., Moleón, M., Gibert, J. P., Botella, F., Mateo-Tomás, P., Olea, P. P., … Sánchez-Zapata, J. A. (2016). Nested species-rich networks of scavenging vertebrates support high levels of interspecific competition. Ecology, 97(1), 95–105. doi:10.1890/15-0212.1

Skinner, J. D., & Chimimba, C. T. (2005). The Mammals of the Southern African Region (Third Edit). Cambridge (UK): Cambridge University Press.

Sobral, M., Silvius, K. M., Overman, H., Oliveira, L. F. B., Raab, T. K., & Fragoso, J. M. V. (2017). Mammal diversity influences the carbon cycle through trophic interactions in the Amazon. Nature Ecology and Evolution, 1, 1670–1676. doi:10.1038/s41559-017-0334-0

Tilman, D., Knops, J., Wedin, D., Reich, P., Ritchie, M., & Siemann, E. (1997). The influence of functional diversity and composition on ecosystem processes. Science, 277(5330), 1300–1302. doi:10.1126/science.277.5330.1300

Villéger, S., Mason, N. W. H., & Mouillot, D. (2008). New multidimensional functional diversity indices for a multifaceted framework in functional ecology. Ecology, 89(8), 2290–2301. doi:10.1890/07-1206.1

Vucetich, J. A., Peterson, R. O., & Waite, T. A. (2004). Raven scavenging favours group foraging in wolves. Animal Behaviour, 67(6), 1117–1126. doi:10.1016/j.anbehav.2003.06.018

Wilson, D. E., & Mittermeier, R. A. (2009). Handbook of the Mammals of the World. Vol. 1. Carnivores. Barcelona (Spain): Lynx Edicions.

Wilson, D. E., & Mittermeier, R. A. (2011). Handbook of the Mammals of the World. Vol. 2. Hoofed Mammals. Barcelona (Spain): Lynx Edicions.

Winfree, R., Fox, J. W., Williams, N. M., Reilly, J. R., & Cariveau, D. P. (2015). Abundance of common species, not species richness, drives delivery of a real-world ecosystem service. Ecology Letters, 18(7), 626–635. doi:10.1111/ele.12424

Zuur, A. F., Ieno, E. N., & Elphick, C. S. (2009). A protocol for data exploration to avoid common statistical problems. Methods in Ecology and Evolution, 1(1), 3–14. doi:10.1111/j.2041-210x.2009.00001.x

