## Supplementary material for "Large home range scavengers support higher rates of carcass removal": File including supplementary tables and figures

Table S1. Summary of the main characteristics of the vertebrate scavenger communities considered, including the total number (N) of carcasses monitored per study area and vertebrate scavenger richness. The table also shows the total abundance of scavenging individuals recorded at each study area, carrion consumption rate and carcass weight (median value and the range of first and third quartiles, Q1-Q3) per monitored carcass.

| | Area | N carcasses | N scavenger species | Abundance per carcass (median; Q1-Q3) | Carrion consumption rate (median kg $h^{-1}$; Q1-Q3) | Carcass weight (median kg; Q1-Q3) |
|---|---|---|---|---|---|---|
| Spain | CC | 72 | 17 | 9.5 (5-19.25) | 0.41 (0.18-1.52) | 15.0 (7.60-31.25) |
| | CZ | 31 | 12 | 33 (14-46.5) | 1 (0.31-1.96) | 33.0 (24.0-60.0) |
| | ES | 11 | 7 | 5 (3-8) | 0.33 (0.21-0.33) | 40.0 (34.5-47.5) |
| South Africa | HiP | 16 | 10 | 18.5 (4-28.5) | 0.83 (0.59-2.26) | 77.5 (55.0-162.5) |
| | Mkhuze | 19 | 14 | 9 (4-34) | 0.61 (0.39-1.39) | 55.0 (44.0-55.0) |

Table S2. Spearman correlation values (ρ) between species abundances at each carcass and carrion consumption rates for the five different areas. ES: Sierra Espuña, CZ: Sierra de Cazorla, CC: Cordillera Cantábrica, MK: Mkhuze, HiP: Hluhluwe-iMfolozi.

| Functional group | Species | ES | CZ | CC | MK | HiP |
|---|---|---|---|---|---|---|
| **Corvids** | *Corvus corax* | -0.07 | -0.02 | -0.04 | | |
| | *Corvus corone* | | -0.04 | 0.06 | | |
| | *Garrulus glandarius* | | -0.20 | | | |
| | *Pica pica* | -0.01 | 0.08 | -0.02 | | |
| | *Corvus albus* | | | | -0.34 | 0.14 |
| **Other raptors** | *Aquila chrysaetos* | 0.12 | 0.28 | 0.14 | | |
| | *Aquila rapax* | | | | -0.06 | -0.03 |
| | *Buteo buteo* | | | -0.16 | | |
| | *Milvus aegyptius* | | | | | 0.37 |
| | *Milvus milvus* | | -0.24 | -0.19 | | |
| **Vultures** | *Aegypius monachus* | | -0.02 | 0.09 | | |
| | *Gypaetus barbatus* | | 0.31 | | | |
| | *Gyps africanus* | | | | -0.25 | 0.23 |
| | *Gyps coprotheres* | | | | 0.26 | |
| | *Gyps fulvus* | | 0.63 | 0.50 | | |
| | *Necrosyrtes monachus* | | | | | 0.08 |
| | *Neophron percnopterus* | | | -0.12 | | |
| | *Torgos tracheliotos* | | | | -0.29 | 0.13 |
| | *Trigonoceps occipitalis* | | | | -0.08 | 0.34 |
| **Omnivores** | *Apodemus* spp. | | | | | |
| | *Potamochoerus larvatus* | | -0.60 | -0.18 | -0.36 | |
| | *Sus scrofa* | 0.02 | | -0.16 | | |
| **Meso-carnivores** | *Canis adustus* | | | | 0.24 | |
| | *Canis familiaris* | -0.51 | -0.32 | -0.28 | | |
| | *Canis mesomelas* | | | | -0.04 | |
| | *Genetta genetta* | | | -0.17 | | |
| | *Genetta tigrina* | | | | -0.47 | |
| | *Ichneumia albicauda* | | | | -0.26 | |
| | *Martes spp* | -0.04 | | -0.24 | | |
| | *Vulpes vulpes* | 0.41 | -0.31 | -0.25 | | |
| **Large-carnivores** | *Canis lupus* | | | 0.11 | | |
| | *Crocuta crocuta* | | | | 0.17 | 0.03 |
| | *Lycaon pictus* | | | | 0.09 | |
| | *Panthera leo* | | | | | 0.2 |
| | *Panthera pardus* | | | | 0.21 | -0.03 |
| | *Ursus arctos* | | | 0.04 | | |

Table S3. Spearman correlation values ($\rho$) between scavenger community-level metrics and carrion consumption rates for the five different areas. ES: Sierra Espuña, CZ: Sierra de Cazorla, CC: Cordillera Cantábrica, MK: Mkhuze, HiP: Hluhluwe-iMfolozi.

| Community metric | ES | CZ | CC | MK | HiP |
|---|---|---|---|---|---|
| **Corvid biomass** | -0.06 | -0.07 | -0.03 | -0.34 | 0.14 |
| **Other raptor biomass** | 0.12 | 0.18 | -0.03 | -0.06 | 0.15 |
| **Vulture biomass** | | 0.64 | 0.50 | -0.24 | 0.26 |
| **Omnivore biomass** | 0.02 | -0.60 | -0.22 | -0.36 | |
| **Meso-carnivore biomass** | 0.03 | -0.45 | -0.40 | -0.19 | |
| **Large-carnivore biomass** | | | 0.12 | 0.21 | 0.30 |
| **Home range** | 0.12 | 0.67 | 0.53 | 0.47 | -0.16 |
| **% obligate scavengers** | | 0.69 | 0.51 | -0.22 | 0.28 |
| **% large carnivorous** | 0.19 | 0.25 | 0.23 | 0.52 | -0.28 |
| **% carnivorous** | 0.19 | 0.71 | 0.53 | 0.37 | -0.14 |
| **% social foraging** | | 0.69 | 0.51 | -0.17 | 0.25 |
| **Functional richness** | -0.03 | -0.01 | 0.05 | -0.31 | 0.29 |
| **Functional dispersion** | 0.05 | -0.22 | -0.04 | -0.13 | 0.15 |
| **Species richness** | -0.10 | -0.08 | -0.12 | -0.25 | 0.40 |
| **Abundance** | -0.03 | 0.60 | 0.37 | -0.34 | 0.38 |
| **Biomass** | -0.12 | 0.19 | 0.26 | -0.24 | 0.41 |

Table S4. Models within the 95% cumulative Akaike weight range relating carrion consumption rate with study area and selected scavenger community attributes: home range, proportion of large carnivorous (%large carnivorous), functional dispersion (FDis), scavenger species richness and abundance. Carcass weight was also included as predictor. Regression coefficients, goodness-of-fit ($r^2$), degrees of freedom (df), AICc and ΔAICc values, and model weight are also shown. +: area / interaction terms were included into the model. Models were fitted using Maximum-Likelihood method.

| **Ranking** | **Intercept** | **Study area** | **Home range** | **% Large carnivorous** | **FDis** | **Scavenger richness** | **Scavenger abundance** | **Study area x species richness** | **Carcass weight** | $r^2$ | **df** | **AICc** | **ΔAICc** | **weight** |
|---|---|---|---|---|---|---|---|---|---|---|---|---|---|---|
| **1** | -0.53 | | 0.74 | | | -0.14 | | | 0.45 | 0.481 | 9 | 483.02 | 0.00 | 0.17 |
| **2** | -0.53 | | 0.70 | | | | | | 0.46 | 0.472 | 8 | 483.36 | 0.34 | 0.14 |
| **3** | -0.54 | | 0.78 | | 0.16 | -0.27 | | | 0.42 | 0.488 | 10 | 483.39 | 0.37 | 0.14 |
| **4** | -0.52 | | 0.70 | | | -0.19 | 0.09 | | 0.44 | 0.483 | 10 | 484.79 | 1.77 | 0.07 |
| **5** | -0.52 | | 0.74 | -0.04 | | -0.15 | | | 0.48 | 0.482 | 10 | 485.15 | 2.13 | 0.06 |
| **6** | -0.53 | | 0.80 | -0.06 | 0.18 | -0.29 | | | 0.46 | 0.489 | 11 | 485.22 | 2.20 | 0.06 |
| **7** | -0.54 | | 0.75 | | 0.15 | -0.29 | 0.07 | | 0.41 | 0.489 | 11 | 485.40 | 2.39 | 0.05 |
| **8** | -0.53 | | 0.72 | | | | -0.04 | | 0.46 | 0.472 | 9 | 485.50 | 2.48 | 0.05 |
| **9** | -0.52 | | 0.71 | -0.02 | | | | | 0.48 | 0.472 | 9 | 485.57 | 2.55 | 0.05 |
| **10** | -0.53 | | 0.70 | | -0.02 | | | | 0.46 | 0.472 | 9 | 485.59 | 2.57 | 0.05 |
| **11** | -0.67 | + | 0.72 | | | -0.08 | | + | 0.60 | 0.532 | 17 | 487.00 | 3.98 | 0.02 |
| **12** | -0.52 | | 0.70 | 0.00 | | -0.19 | 0.09 | | 0.44 | 0.483 | 11 | 487.12 | 4.10 | 0.02 |
| **13** | -0.68 | + | 0.76 | | 0.18 | -0.16 | | + | 0.60 | 0.539 | 18 | 487.27 | 4.25 | 0.02 |
| **14** | -0.53 | | 0.78 | -0.05 | 0.17 | -0.29 | 0.02 | | 0.45 | 0.489 | 12 | 487.56 | 4.54 | 0.02 |
| **15** | -0.51 | | 0.75 | -0.05 | | | -0.07 | | 0.50 | 0.473 | 10 | 487.57 | 4.55 | 0.02 |
| **16** | -0.53 | | 0.72 | | 0.00 | | -0.04 | | 0.46 | 0.472 | 10 | 487.80 | 4.78 | 0.02 |
| **17** | -0.52 | | 0.71 | -0.02 | -0.01 | | | | 0.48 | 0.472 | 10 | 487.84 | 4.83 | 0.02 |
| **18** | -0.76 | + | 0.73 | | 0.18 | -0.31 | | | 0.56 | 0.503 | 14 | 488.47 | 5.45 | 0.01 |

| | | | | | | | | | | | | | |
|---|---|---|---|---|---|---|---|---|---|---|---|---|---|
| **19** | -0.73 | + | 0.68 | | -0.17 | | | 0.57 | 0.494 | 13 | 488.52 | 5.51 | 0.01 |
| **20** | -0.65 | + | 0.68 | | -0.10 | 0.11 | + | 0.59 | 0.534 | 18 | 488.86 | 5.84 | 0.01 |
| **21** | -0.66 | + | 0.72 | 0.17 | -0.18 | 0.10 | + | 0.59 | 0.541 | 19 | 489.29 | 6.27 | 0.01 |

Fig. S1. Variation in functional group biomass across the five areas. ES: Sierra Espuña, CZ: Sierra de Cazorla, CC: Cordillera Cantábrica, MK: Mkhuze, HiM: Hluhluwe-iMfolozi. Study areas are represented by different symbols depending on their scavenger composition: wild board: only facultative scavengers (ES); vulture: facultative + obligate (CZ and MK); lion: facultative + obligate + top predators (CC and HiP).

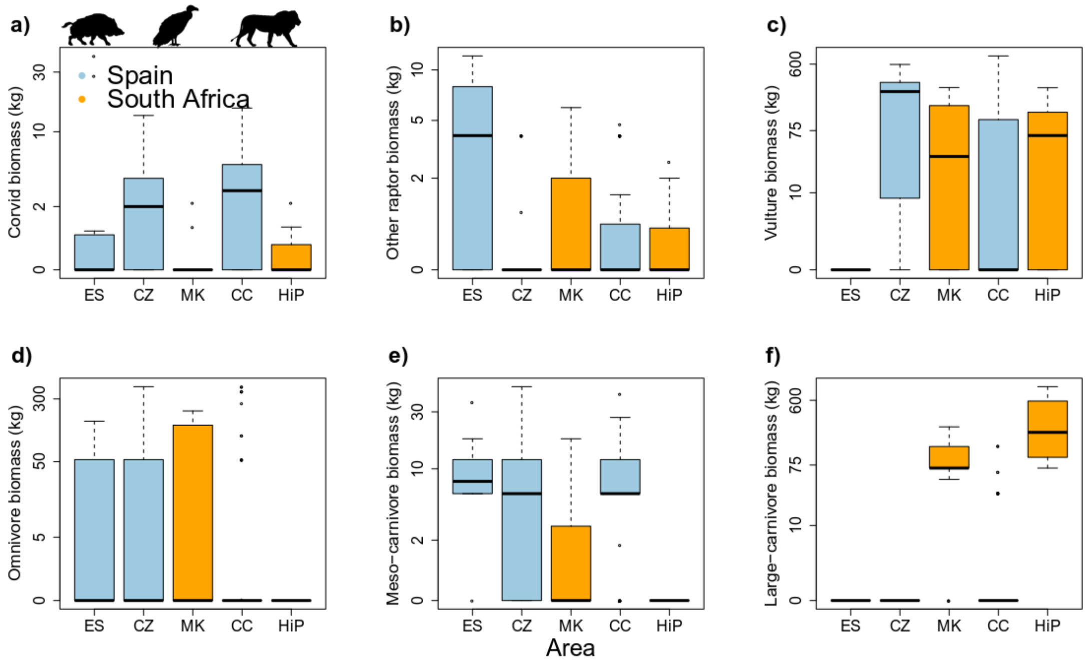

Fig S2. Boxplots showing variation in species richness, community abundance, community biomass, functional richness, functional dispersion and consumption rate across landscapes. ES: Sierra Espuña, CZ: Sierra de Cazorla, CC: Cordillera Cantábrica, MK: Mkhuze, HiM: Hluhluwe-iMfolozi. Study areas are represented by different symbols depending on their scavenger composition: wild board: only facultative scavengers (ES); vulture: facultative + obligate (CZ and MK); lion: facultative + obligate + top predators (CC and HiP).

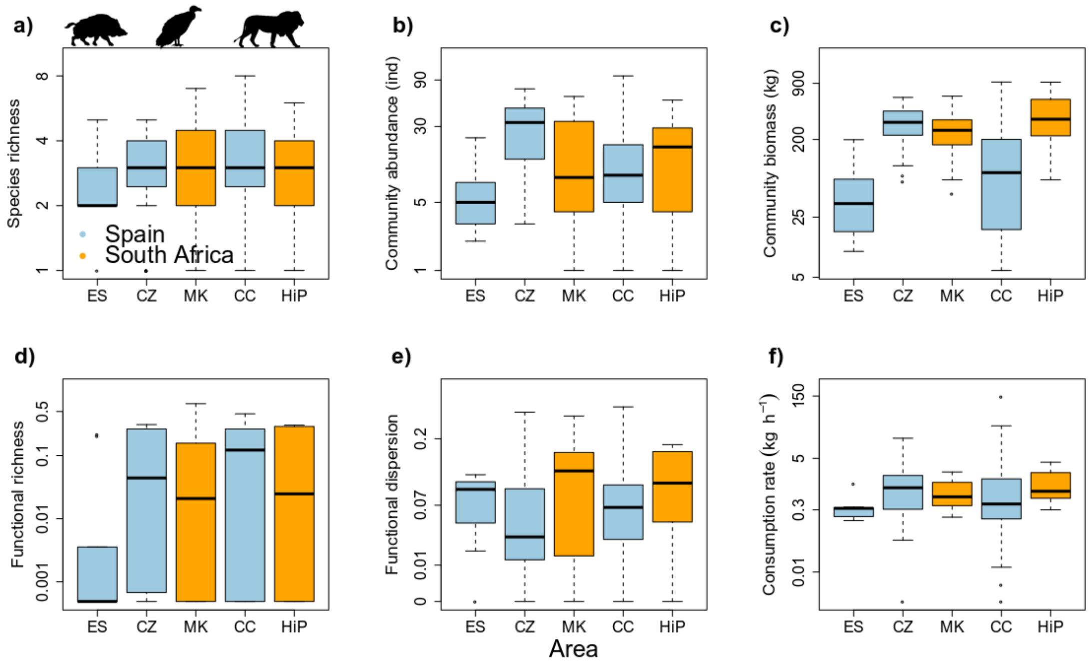

Fig. S3. Mean (± 95% CI) of spearman correlations between carrion consumption rates and species abundances at each for the five study areas. Species results are shown using different symbols for each region (square: Spain, triangle: South Africa). Species with less than five occurrences within a study area are not shown. See Table S2 for more details.

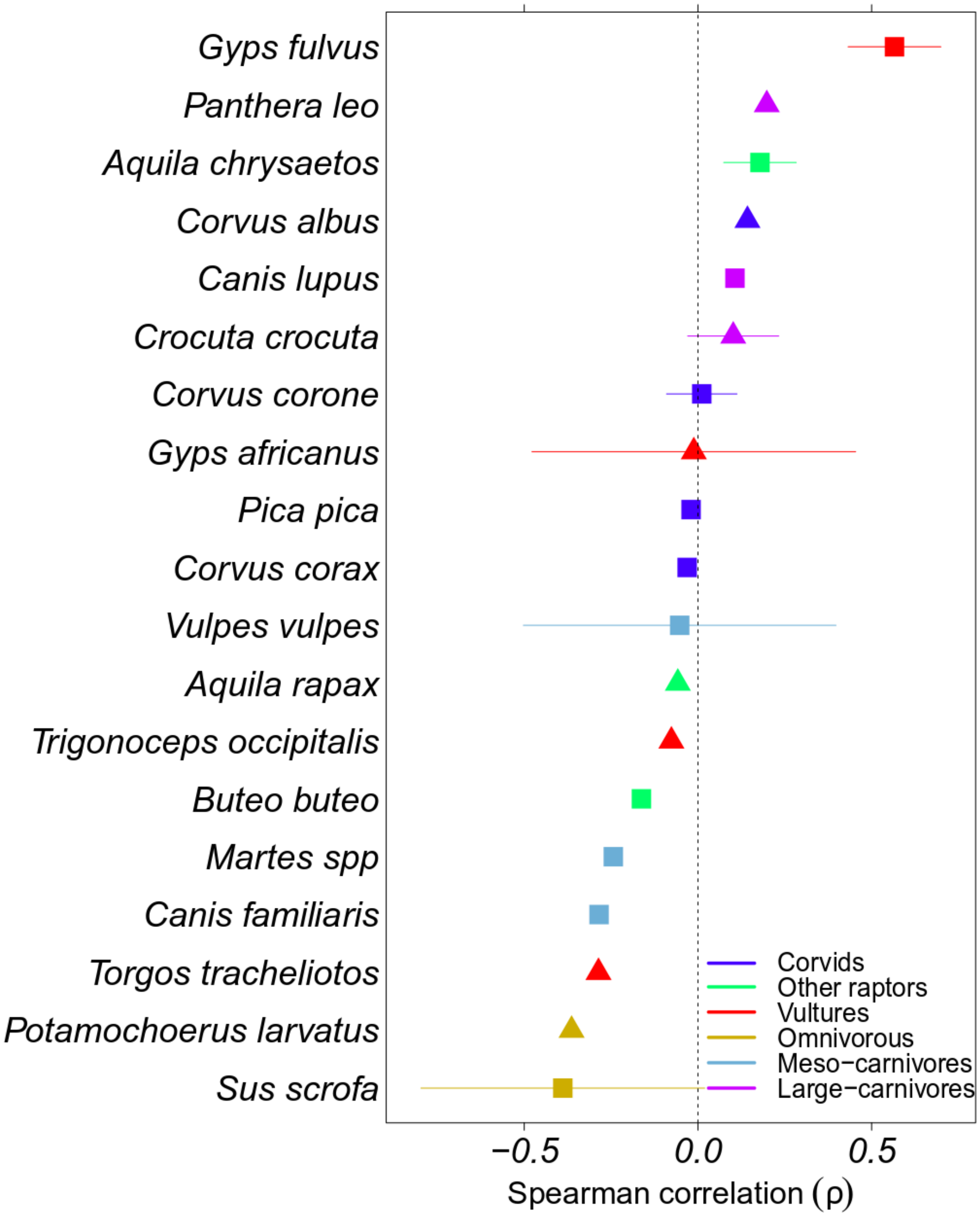
