## Supplementary material for "Large home range scavengers support higher rates of carcass removal": Files including R functions and code to reproduce some of our statistical analyses: 0_FD_functions.R

```
#################################################################################################################
#                                                                                                               #
# Supplementary Appendix S1                                                                                     #
#                                                                                                               #
# R script to generate trait-based metrics                                                                      # 
#                                                                                                               #
#                                                                                                               #
# Large home range scavengers support higher rates of carcass removal                                           #
#                                                                                                               #
# Gutiérrez-Cánovas, C., Moleón, M., Mateo-Tomás, P., Olea, P.P., Sebastián-González, E.                        #
# Sánchez-Zapata, J.A.                                                                                          #
#                                                                                                               #
# Code written by Cayetano Gutiérrez-Cánovas,                                             #
#                                                                                                               #
#################################################################################################################

# fric_3d() estimates the Functional Richness of a set of communties
# This function computes the hypervolume to estimate how each community fills
# the functional space
#
# Inputs:
# taxa: community data
# fpc: functional space
# m: number of axes to select
# prec: convex hull precision ("Qt" or "QJ")
#
# Output:
# a vector with the Functional Richness of each community

fric_3d<-function(taxa,fpc,m,prec=c("Qt","QJ")){
  fric.3d<-rep(NA,nrow(taxa))
  convhulln(fpc[,1:m], c("FA",prec))$vol->fric.3d.max
  specnumber(taxa)->ric
  for (com in 1:nrow(taxa)){
    fpc[which(unlist(rep(taxa[com,]))>0),1:m]->tr.com
    if (ric[com]>=m+1) convhulln(tr.com, c("FA",prec))$vol/fric.3d.max->fric.3d[com] else NA->fric.3d[com]
  }
  return(fric.3d)
}

# calc.FR() estimates the Functional Redundancy (FR) of a set of communties
# This function estimates the taxonomic richness for each taxonomic group,
# estating FR as the ratio between species richness and the number of functional groups
# in a given community.
#
# Inputs:
# taxa: community data
# groups: a grouping vector with the Funtional Groups for each taxon
#
# Outputs:
#
# $nbsp: taxonomic richness
# $ric.fgrs: the taxonomic richness for each Functional Group
# $sd.fgr: the among-Functional-group SD for taxonomic richness for each community
# $FGR: the number of Functional Gruoups for each community
# $FR: the Functional Richness for each community

calc.FR<-function(taxa,groups){
  list()->res
  
  if (ncol(taxa)!=length(groups)) stop("Trait and taxonomic matrix have different number of taxa")
  unique(groups)->gr.names
  specnumber(taxa)->res$nbsp

  res$ab.fgrs<-res$ric.fgrs<-data.frame(matrix(NA,nrow(taxa),length(unique(groups))))
  colnames(res$ab.fgrs)<-colnames(res$ric.fgrs)<-paste("FR",gr.names,sep="")
  
  j<-0
  
  for (i in gr.names) {
    j<-j+1
    if(is.vector(taxa[,which(groups==i)])==T) decostand(taxa[,which(groups==i)],"pa")->res$ric.fgrs[,j] else specnumber(taxa[,which(groups==i)])->res$ric.fgrs[,j]
    }
  
  j<-0
  
  for (i in gr.names) {
  j<-j+1
    if(is.vector(taxa[,which(groups==i)])==T) taxa[,which(groups==i)]->res$ab.fgrs[,j] else rowSums(taxa[,which(groups==i)])->res$ab.fgrs[,j]
  }
  
  apply(res$ric.fgrs,1,sd)->res$sd.fgr
  specnumber(res$ric.fgrs)->res$FGR # functional group richness estimation
  res$FR<-res$nbsp/res$FGR
  res$FR.ab<-rowSums(taxa)/res$FGR
  return(res)
}

# fdisp_k() estimates the Functional Dispersion of a set of communties
# This function computes the weighted mean distance to community centroid in
# the functional space
# 
# Function modified from Laliberté & Legendre (2010) Ecology
#
# Inputs:
# d: trait dissimilarity matrix
# a: community data
# m: number of axes to select
# tol: tolerance threshold to test whether the distance matrix is Euclidean
#
# Output:
# FDis: a vector with the Functional Dispersion of each community
# eig: eigenvectors of each functional axis
# vectors: functional axes

fdisp_k<-function (d, a, m, tol = 1e-07) 
{
  if (!inherits(d, "dist")) 
    stop("'d' must be a 'dist' object.")
  n <- attr(d, "Size")
  if (is.null(attr(d, "Labels"))) 
    stop("'d' must have labels.", "\n")
  else sn.d <- attr(d, "Labels")
  if (missing(a)) {
    ab.names <- list("Community1", sn.d)
    a <- matrix(1, 1, n, dimnames = ab.names)
  }
  com <- nrow(a)
  if (ncol(a) != n) 
    stop("Number of columns in 'a' must be equal to the number of objects in 'd'.")
  if (is.null(colnames(a))) 
    stop("'a' must have column names", "\n")
  else sn.a <- colnames(a)
  if (any(sn.d != sn.a)) 
    stop("Species labels in 'd' and 'a' need to be identical and ordered alphabetically (or simply in the same order).", 
         "\n")
  a[which(is.na(a))] <- 0
  abun.sum <- apply(a, 1, sum)
  if (any(abun.sum == 0)) 
    warning("At least one community has zero-sum abundances (no species).", 
            "\n")
  abun.sum2 <- apply(a, 2, sum)
  if (any(abun.sum2 == 0)) 
    warning("At least one species does not occur in any community (zero total abundance across all communities).", 
            "\n")
  if (any(is.na(d))) 
    stop("NA's in the distance matrix.", "\n")
  A <- matrix(0, ncol = n, nrow = n)
  A[row(A) > col(A)] <- -0.5 * d^2
  A <- A + t(A)
  G <- bicenter.wt(A)
  e <- eigen(G, symmetric = TRUE)
  vectors <- e$vectors
  eig <- e$values
  w0 <- eig[n]/eig[1]
  if (w0 > -tol) 
    r <- sum(eig > (eig[1] * tol))
  else r <- length(eig)
  vectors <- vectors[, 1:r, drop = FALSE] %*% diag(sqrt(abs(eig <- eig[1:r])), 
                                                   r)
  dimnames(vectors) <- list(colnames(a), NULL)
  pos <- eig > 0
  if (m>0) pos<-c(pos[1:m],rep(F,length(pos)-m))
  
  avg.dist.cent <- rep(NA, nrow(a))
  names(avg.dist.cent) <- row.names(a)
  for (i in 1:com) {
    pres <- which(a[i, ] > 0)
    nb.sp <- nrow((unique(vec <- vectors[pres, , drop = F])))
    if (nb.sp >= 2) {
      w <- a[i, pres]
      centroid <- apply(vec, 2, weighted.mean, w = w)
      dist.pos <- sweep(vec[, pos, drop = F], 2, centroid[pos])
      dist.pos <- rowSums(dist.pos^2)
      if (any(!pos)) {
        dist.neg <- sweep(vec[, !pos, drop = F], 2, centroid[!pos])
        dist.neg <- rowSums(dist.neg^2)
      }
      else dist.neg <- 0
      zij <- sqrt(abs(dist.pos - dist.neg))
      avg.dist.cent[i] <- weighted.mean(zij, w)
    }
    else avg.dist.cent[i] <- 0
  }
  return(list(FDis = avg.dist.cent, eig = eig, vectors = vectors))
}

## Run the following lines. These introduce post-hoc methods for 'gls' objects.

# Source: http://rstudio-pubs-static.s3.amazonaws.com/13472_0daab9a778f24d3dbf38d808952455ce.html
model.matrix.gls <- function(object, ...) {
  model.matrix(terms(object), data = getData(object), ...)
}
model.frame.gls <- function(object, ...) {
  model.frame(formula(object), data = getData(object), ...)
}
terms.gls <- function(object, ...) {
  terms(model.frame(object), ...)}
x.quasipoisson <- function(...)
{
  res <- quasipoisson(...)
  res$aic <- poisson(...)$aic
  res
}

dfun <- function(object) {
  with(object,sum((weights * residuals^2)[weights > 0])/df.residual)
}

#  This is important to back-standardise data in plots
back_st<-function(x,x_mean,x_sd){return(x*x_sd+x_mean)}
```
