## Supplementary material for "Large home range scavengers support higher rates of carcass removal": Files including R functions and code to reproduce some of our statistical analyses: 0_quality_funct_space.R

```
#################################################################################################################################
## quality_funct_space: R function for computing the quality of functional multidimensional functional spaces                  ##
##                                                                                                                             ##
##                  Given a functional matrix, the function computes the quality (i.e. mean squared-deviation between          ##
##                  initial functional distance and standardized distance in the functional space) for all the                 ##
##                  multidimensional functional spaces from 2 to N dimensions (N selected by the user).                        ## 
##                                                                                                                             ##
##  Original code by Eva Maire & Sébastien Villéger in:                                    ##   
##                                                                                                                             ##
##    Maire, E., Grenouillet, G., Brosse, S., & Villéger, S. (2015) How many dimensions are needed to                          ##
##    accurately assess functional diversity? A pragmatic approach for assessing the quality of functional                     ##
##    spaces. Global Ecology and Biogeography, 24, 728–740.                                                                    ##
##                                                                                                                             ##
##                                                                                                                             ##
##  Code modified and adapted by Cayetano Gutiérrez-Cánovas                                  ##
##                                                                                                                             ##
##                                                                                                                             ##
##     INPUTS:                                                                                                                 ##
##                                                                                                                             ##
##      - "mat_funct" : a species x traits functional matrix (NA are not allowed).                                             ##
##                        Traits could be of different types (e.g. numeric, ordinal, nominal)                                  ##
##                                                                                                                             ##
##      - "nbdim" : maximum number of dimensions for multidimensional functional spaces. By default, nbdim=7                   ##
##      		Final number of dimensions depends on the number of positive eigenvalues (after correction) obtained with PCoA     ##
##                                                                                                                             ##
##      - "metric" : metric to be used to compute functional distance, "Euclidean" or "Gower" (=default)                       ##
##                                                                                                                             ##
##      - "plot" : character string to set the name of the jpeg file for plots illustrating the quality of functional spaces   ##
##                  NA means no plot                                                                                           ##
##                                                                                                                             ##
##     NB: 1/ high value for 'nbdim' can increase computation time                                                             ##
##         2/ if at least one trait is not numeric, 'metric' should equal 'gower'                                              ##
##         3/ if metric=Euclidean, functional traits are scaled (mean=0, sd=1) before computing functional distances           ##
##         4/ R libraries 'ape', 'clue', 'cluster', 'FD', 'geometry', 'gtools' are required                                    ##
##                                                                                                                             ##
##                                                                                                                             ##
##      OUTPUTS: a list with                                                                                                   ##
##                                                                                                                             ##
##      - $ meanSD : a vector with mean squared deviation values for all the functional spaces tested							             ##
##            names are 'm_kD' (with k=2:nbdim) for multidimensional spaces                                                    ##
##                                                                                                                             ##
##      - $ mat_dissim : matrix of functional dissimilarities between species based on their traits values                     ##
##                                                                                                                             ##
##      - $ fpc : coordinates of species in the nbdim multidimensional functional space (PCoA)                                 ##
##                                                                                                                             ##
##                                                                                                                             ##
##                                                                                                                             ##
#################################################################################################################################

quality_funct_space <- function(mat_funct, nbdim=7, metric="Gower",corr_method=c("none","cailliez","lingoes")) {

#loading required libraries
require(ape)
require(clue)
require(cluster)
require(FD)
require(geometry)
require(gtools)

################################################################################################################################# 

# checking data
if (sum(is.na(mat_funct))!=0)   {  warning(" NA detected in 'funct_mat' ")     }
if (nbdim<2)   {  stop(" 'nbdim' must be higher than 1")     }
if (nrow(mat_funct)<3)   {  stop(" there must be at least 3 species in 'funct_mat' ")     }
if (metric=="Euclidean" & sum(apply(mat_funct,2,is.numeric))!=ncol(mat_funct) )  { stop("using Euclidean distance requires that all traits are continuous")   }
if (metric=="Euclidean" & nbdim>ncol(mat_funct) )  { stop("using Euclidean distance requires less dimensions than number of traits")   }
if (metric=="Gower" & nbdim>nrow(mat_funct) )  { stop("using Gower's distance requires less dimensions than number of species")   }

  
# computing functional dissimilarity between species given their traits values
if (metric=="Gower") mat_dissim<-gowdis(mat_funct,ord="classic")
if (metric=="Euclidean") mat_dissim<-dist(scale(mat_funct)) # scaling if continuous traits

# lists to store distnaces matrices
dist_raw<-list()
dist_st<-list()
  
################################
# computing PCoA using Caillez correction
mat_pcoa<-pcoa(mat_dissim , correction=corr_method)

# changing number of dimensions given number of positive eigenvalues
nbdim<-min(nbdim,ncol(mat_pcoa$vectors) )

# keeping species coordoinates on the 'nbdim' axes
mat_coord<-mat_pcoa$vectors[,1:nbdim]
row.names(mat_coord)<-row.names(mat_funct)
colnames(mat_coord)<-paste("PC",1:nbdim,sep="")

# computing Euclidean distances between species in the (nbdim-1) multidimensionnal functional spaces 
for (k in 2:nbdim) {
    eval(parse(text=paste("dist_",k,"D<-dist(mat_coord[,1:",k,"],method='euclidean')", sep="")))
    eval(parse(text=paste("dist_raw$m_",k,"D<-dist_",k,"D", sep="")))
  } # end of k
    
################################
# computing mean squared deviation between initial distance and standardized final distance in the functional space
meanSD<-rep(NA,nbdim-1) ; names(meanSD)<-c(paste(paste("m_",2:nbdim,"D",sep="")))

x<-mat_dissim # initial distance
S<-nrow(mat_funct) # species richness

# for muldimensionnal spaces
for (k in 2:nbdim)  {
eval(parse(text=paste("y<-dist_",k,"D",sep="")))
yst<- y/max(y) * max(x)
eval(parse(text=paste("dist_st$m_",k,"D<-dist_",k,"D", sep="")))
meanSD[paste("m_",k,"D",sep="")]<-round( ( (sum((x-yst)^2)) / (S*(S-1)/2) ) ,6)
}  # end of k

# list of outputs
res<-list(meanSD=meanSD, mat_dissim=mat_dissim, fpc=mat_coord)
     
invisible(res)

} # end of function quality_funct_space

################################################################################################################################
################################################################################################################################
```
