## Supplementary material for "Large home range scavengers support higher rates of carcass removal": Files including R functions and code to reproduce some of our statistical analyses: 1_biometric estimation.R

```
#################################################################################################################
#                                                                                                               #
# Supplementary Appendix S1                                                                                     #
#                                                                                                               #
# R script to estimate scavenger community metrics (functional identity, functional diversity,                  #
# functional equivalence)                                                                                       # 
#                                                                                                               #
#                                                                                                               #
# Large home range scavengers support higher rates of carcass removal                                           #
#                                                                                                               #
# Gutiérrez-Cánovas, C., Moleón, M., Mateo-Tomás, P., Olea, P.P., Sebastián-González, E.                        #
# Sánchez-Zapata, J.A.                                                                                          #
#                                                                                                               #
# Code written by Cayetano Gutiérrez-Cánovas,                                             #
#                                                                                                               #
#################################################################################################################

setwd("c:/Users/Tano/Dropbox/Trabajos en curso/scavenging/analysis/def")
setwd("your_folder")
plot_folder<-paste(getwd(),"/plots/",sep="")

# Loading libraries
library(FD)
library(vegan)
library(car)
library(multcomp)
library(ade4)
library(ape)
library(plyr)
library(sqldf)

#loading additional functions
source("0_FD_functions.R")
source("0_quality_funct_space.R")

### Loading data
taxa<-read.table("taxa_ab.txt",h=T,sep="\t")
tr<-read.table("traits.txt",h=T,sep="\t")
env<-read.table("env.txt",h=T,sep="\t")

# Seleccting carcasses
env<-sqldf("select * from env,taxa where env.carcass = taxa.carcass")[,1:9]

# Assigning rownames and removing labels to analyise data
rownames(taxa)<-taxa[,1]
taxa<-(taxa[,-c(1)])

rownames(tr)<-tr[,1]
tr<-(tr[,-1])

rownames(env)<-env[,1]
env<-(env[,-c(1:2)])

# Checking species and row order
any(rownames(tr)!=colnames(taxa))
any(rownames(taxa)!=rownames(env))

# Are there non-occurring species?
any(colSums(taxa)==0) 

# Are there sites with no species?
any(rowSums(taxa)==0) 

# Creating consumption_rate vector
c.rate<-log(env$cons_rate)

# Estimating biomass for each species
t(t(taxa) * tr$body_mass)->taxa_bio

# Assessing skewness

# Checking the distribution of the quantitative traits

# Histograms of quantitative variables
par(mfrow=c(2,2))
hist(tr$range)
hist(tr$body_mass)
hist(tr$fecundity)
hist(tr$longevity)

# Histograms of transformed quantitative variables
hist(log(tr$range))
hist(log(tr$body_mass))
hist(log(tr$fecundity))
hist(sqrt(tr$longevity))

# Transformations
log(tr$body_mass)->tr$body_mass
log(tr$fecundity)->tr$fecundity

# ordering ordinal traits and areas

tr$predator<-ordered(tr$predator,levels=c("non_pred","meso_pred","top_pred"))
tr$social<-ordered(tr$social,levels=c("solitary","group","social"))

env$type_reg<-ordered(env$type_reg,levels=c("F-M","VF-M","VF-S","PVF-M","PVF-S"))

###################################################################################################
# Functional space and groups
################################################################################################### 

# calculate species x species Gower dissimilarity matrix based on functional traits
gowdis(tr[-c(14:15)])->tr.dist

# Estimating the optimum number of dimensions
qual_fs<-quality_funct_space(tr, nbdim=10, metric="Gower","none")
qual_fs$meanSD<0.01 # 2D seems to be an appropiate number of dimensions
qual_fs$meanSD

# checking for negative eigenvalues
dudi.pco(tr.dist,scannf = F,nf=5)->tr.pco
length(which(tr.pco$eig<0)) # No negative eigenvalues licenced us to use Ward clustering method
tr.pco$eig[1:3]/sum(tr.pco$eig)
sum(tr.pco$eig[1:2]/sum(tr.pco$eig)) # 79.7%


# Correlation between axes and original quantiative variables
cor(tr[,which(sapply(tr,is.numeric))],tr.pco$li)

for (i in 1:5) {print(i);print(summary(aov(tr.pco$li[,i]~tr$mobility)))} # axis 1
for (i in 1:5) {print(i);print(summary(aov(tr.pco$li[,i]~tr$social)))} # axis 3, 1, 4
for (i in 1:5) {print(i);print(summary(aov(tr.pco$li[,i]~tr$migrant)))} # axis 5, 3
for (i in 1:5) {print(i);print(summary(aov(tr.pco$li[,i]~tr$scavenger)))} # axis 1, 5
for (i in 1:5) {print(i);print(summary(aov(tr.pco$li[,i]~tr$predator)))} # axis 2, 5, 1
for (i in 1:5) {print(i);print(summary(aov(tr.pco$li[,i]~tr$diet)))} # axis 2, 1
for (i in 1:5) {print(i);print(summary(aov(tr.pco$li[,i]~tr$activity)))} # axis 1, 5
for (i in 1:5) {print(i);print(summary(aov(tr.pco$li[,i]~tr$sight)))} # axis 1, 2, 5
for (i in 1:5) {print(i);print(summary(aov(tr.pco$li[,i]~tr$smell)))} # axis 1

par(mfrow=c(4,3))
boxplot(tr.pco$li[,1]~tr$mobility) # Axis 1
boxplot(tr.pco$li[,3]~tr$social) # Axis 3
boxplot(tr.pco$li[,5]~tr$migrant) # Axis 5, 3
boxplot(tr.pco$li[,5]~tr$scavenger) # Axis 1, 5
boxplot(tr.pco$li[,2]~tr$predator) # Axis 2
boxplot(tr.pco$li[,2]~tr$diet) # Axis 1
boxplot(tr.pco$li[,1]~tr$activity) # Axis 1
boxplot(tr.pco$li[,1]~tr$sight) # Axis 1
boxplot(tr.pco$li[,1]~tr$smell) # Axis 1
par(mfrow=c(1,1))

# Two axes are ecologically meaningful
# Axis 1: Positively related with aerial mobility (F=277.0, p<0.001), social feeding (F=4.191, p=0.024), 
# obligate scavenge (F=37.8,p<0.001), non-predator animals (F=7.2, p=0.003), carnivorous diet (F=19.78,p<0.001), 
# carnivorous diet (F=19.8, p<0.001), diurnal activity (F=136.2, p<0.001), high sight (F=10.2,p=0.003), 
# low smell (F=277.8, p<0.001) and negatively with meso and top predation, and group and solitary feeding.
# Axis 2: Positive related with range (R=-0.66), body_mass (R=-0.72), top-predation (F=18.1, p<0.001) and 
# carnivorous diet (F=27.1, p<0.001)

# Classifying species into functional groups
tr.clust <- hclust(tr.dist, method = "ward.D")

# Five groups were chosen
cut.g <- 6
f.gr <- factor(cutree(tr.clust, k = cut.g), labels= c("Vultures", "Omnivores", "Other raptors", "Meso-carnivores", "Large carnivores","Corvids"))

new.fgr<-gsub("n", "", f.gr)
names(new.fgr)<-names(f.gr)
f.gr<-as.factor(new.fgr)

write.table(f.gr, "f_gr.txt", sep="\t")

# Setting group colours
fgr.col<-c("#4600FFFF","#CC00FFFF",blues9[5], "gold3","#00FF66FF","#FF0000FF")

# How groups spread in the functional space
pdf(file=paste(plot_folder,"groups.pdf",sep=""),onefile=T,width=14,height=7,useDingbats=FALSE,)
par(mfrow=c(1,2), mar=c(3,3,3,3))
plot(as.phylo(tr.clust), cex = 1.1, label.offset= 0.05, tip.color = fgr.col[f.gr])
s.class(tr.pco$li[,1:2],fac = f.gr, col=fgr.col,clabel=1.5, cpoint=3,pch=c(15,16,16,16,15,15)[f.gr],cellipse=0)
dev.off()

# FG1 avian, large, obligate scavengers
# FG2 terrestrial,large-size, omnivores
# FG3 large, aerial, meso and predator carnivorous
# FG4 terrestrial,  meso-predators
# FG5 terrestrial, large, top-predators 
# FG6 avian, small, omnivores, meso-predators 

summary(tr[which(f.gr==1),-c(8,9)])
summary(tr[which(f.gr==3),-c(8,9)])
summary(tr[which(f.gr==6),-c(8,9)])

summary(tr[which(f.gr==2),-c(8,9)])
summary(tr[which(f.gr==4),-c(8,9)])
summary(tr[which(f.gr==5),-c(8,9)])

summary(exp(tr[which(f.gr==1),c(8,9)]))
summary(exp(tr[which(f.gr==3),c(8,9)]))
summary(exp(tr[which(f.gr==6),c(8,9)]))

summary(exp(tr[which(f.gr==2),c(8,9)]))
summary(exp(tr[which(f.gr==4),c(8,9)]))
summary(exp(tr[which(f.gr==5),c(8,9)]))

#####################################################
### Scavenger community metrics                   ###
#####################################################

# Funcional identity

calc.FR(taxa_bio,f.gr)->FR.dat # Functional redundancy data

# back-transforming data to estimate the means
exp(tr$body_mass)->tr$body_mass
exp(tr$fecundity)->tr$fecundity
functcomp(tr[,-c(14, 15)],as.matrix(taxa_bio),CWM.type = "all")->cwm

# Functional diversity
fric_3d(taxa, tr.pco$li, m=2, prec="Qt")->FRic # functional richness

FRic[which(is.na(FRic))]<-min(FRic,na.rm=T) # removing NAs

fdisp_k(tr.dist,taxa_bio,m=2)$FDis->FDis # functional dispersion

# Equivalence
specnumber(taxa)->ric # species richness
diversity(taxa)->shannon # shannon
rowSums(taxa)->abun # abundance
rowSums(taxa_bio)->bio # biomass

# Saving results

dat<-data.frame(env,
                FR1=FR.dat$ab.fgrs[,1],
                FR2=FR.dat$ab.fgrs[,2],
                FR3=FR.dat$ab.fgrs[,3],
                FR4=FR.dat$ab.fgrs[,4],
                FR5=FR.dat$ab.fgrs[,5],
                FR6=FR.dat$ab.fgrs[,6],
                FR=FR.dat$FR.ab,
                FRic=FRic, 
                FDis=FDis,
                tax.ric=ric,
                abun=abun,
                bio=bio,
                cwm)


write.table(dat,"dat_final_new.txt",sep="\t")
```
