## Supplementary material for "Large home range scavengers support higher rates of carcass removal": Files including R functions and code to reproduce some of our statistical analyses: 2_descriptive_plots.R

```
#################################################################################################################
#                                                                                                               #
# Supplementary Appendix S1                                                                                     #
#                                                                                                               #
# R script to create descriptive plots and figures                                                              # 
#                                                                                                               #
#                                                                                                               #
# Large home range scavengers support higher rates of carcass removal                                           #
#                                                                                                               #
# Gutiérrez-Cánovas, C., Moleón, M., Mateo-Tomás, P., Olea, P.P., Sebastián-González, E.                        #
# Sánchez-Zapata, J.A.                                                                                          #
#                                                                                                               #
# Code written by Cayetano Gutiérrez-Cánovas,                                             #
#                                                                                                               #
#################################################################################################################

# Working directory
setwd("c:/Users/Tano/Dropbox/Trabajos en curso/scavenging/analysis/def")
setwd("your_folder")
plot_folder<-paste(getwd(),"/plots/",sep="")
assum_folder<-paste(getwd(),"/plots/assum/",sep="")

# Loading libraries
library(car)
library(lattice)
library(usdm)
library(plyr)
library(MuMIn)
library(nlme)

#### Loading data
dat<-read.table("dat_final.txt",h=T,sep="\t")
taxa<-read.table("taxa_ab.txt",h=T,sep="\t")
traits<-read.table("traits.txt",h=T,sep="\t")
f.gr<-read.table("f_gr.txt",h=T,sep="\t")

# Arranging matrices
dat[,1]->rownames(dat)
dat[,-1]->dat

rownames(f.gr)->spp.names

# Arranging functional group matrix
f.gr<-f.gr$x
names(f.gr)<-spp.names

###########################
# Variable transformation
###########################

q<-sapply(dat,class)=="numeric" | sapply(dat,class)=="integer"# selecting quantitative variables

par(mfrow=c(3,3))
for (i in which(q==T)) hist(dat[,i], main=names(dat)[i])
par(mfrow=c(1,1))

log(log(dat$weight_kg))->dat$weight_kg
log(dat$cons_rate)->dat$cons_rate
log(dat$body_mass)->dat$body_mass
hist(car::logit(dat$activity_both/100))

sqrt(dat$FDis)->dat$FDis

log(dat[,c("FR1","FR2","FR3","FR4","FR5","FR6")]+1)->dat[,c("FR1","FR2","FR3","FR4","FR5","FR6")]

log(dat[,c("FR","FRic","tax.ric","abun","bio")])->dat[,c("FR","FRic","tax.ric","abun","bio")]

car::logit(dat[,c("social_social","scavenger_obligate","predator_non_pred","predator_top_pred",
                  "activity_nocturnal","sight_low")])->dat[,c("social_social","scavenger_obligate",
                                                                            "predator_non_pred","predator_top_pred","activity_nocturnal","sight_low")]

par(mfrow=c(3,3))
for (i in which(q==T)) hist(dat[,i], main=names(dat)[i])
par(mfrow=c(1,1))


##################################################################
### Landscape features influencing biodiversity facets
##################################################################

col.reg<-c(blues9[4], blues9[4], "orange", blues9[4], "orange")
landscape<-ordered(dat$type_reg,labels=c("F-M","FO-M","FO-S","FOP-M","FOP-S"), levels=c("F-M","VF-M","VF-S","PVF-M","PVF-S"))

#####################################################
# Boxplots  showing community metrics per study area
#####################################################

sel.var<-c("tax.ric", "abun", "bio", "FRic", "FDis", "cons_rate")

trans<-c(rep("log", 4), "sqrt", "log")

name.var<-c("Species richness", "Community abundance (ind)", "Community biomass (kg)", 
            "Functional richness", "Functional dispersion",expression(Consumption ~ rate ~ (kg~ ~ h^{-1})))

let<-paste(letters,")",sep = "")

var.range<-list(c(1, 2, 4, 8),
                c(1, 5, 30, 90),
                c(5, 25, 200, 900),
                c(0.001, 0.01, 0.10, 0.50),
                c(0, 0.01, 0.07, 0.20),
                c(0.01, 0.3, 5, 150))

pdf(file=paste(plot_folder,"bio_boxplots2.pdf",sep=""),onefile=T,width=12,height=7)

    par(mfrow=c(2,3), cex.lab=1.7, cex.axis=1.5, mar=c(4,5,4,4))
    
    a=0
    
    for (i in sel.var){
      a=a+1
    
        if(a!=4) boxplot(dat[, i] ~ landscape, col=col.reg, cex=0.75, ylab=name.var[a], xaxt="n", yaxt = "n", 
                ylim=c(min(dat[, i]), max(dat[,i])*1.1), xlab="")
      
        if(a==4) boxplot(dat[, i] ~ landscape, col=col.reg, cex=0.75, ylab=name.var[a], xaxt="n", yaxt = "n", 
                         ylim=c(min(dat[, i]), log(1.2)), xlab="")
        
             if (a==1) legend("bottomleft", c("Spain", "South Africa"), col=c(blues9[4], "orange"), pch=16,  bty="n", cex=2.25,
                         y.intersp=0.75)
        mtext(let[a], line = 1.0, adj = -0.2, cex = 1.4, font = 2)
        
        if(a!=4) let.pos<-as.numeric(max(dat[, i])*1.08) else let.pos<-log(0.99)
        
          #text(x=c(1,2,3,4,5),y=let.pos, as.character(bio_res[a, 4:ncol(bio_res)]), cex=1.75)
        
        axis(1, 1:5, c("ES", "CZ", "MK", "CC", "HiP"), cex=1.4)
        
        if (trans[a]=="sqrt") axis(2, sqrt(var.range[[a]]), var.range[[a]])
        if (trans[a]=="log") axis(2, log(var.range[[a]]), var.range[[a]])
        if(a==5) mtext("Area", line= -22.5, cex=1.5)
      }

dev.off()

########################################################################
## Plot showing functional group biomasses per study area
########################################################################

sel.var<-paste("FR", 1:6, sep="")

name.var<-paste(c("Corvid", "Other raptor", "Vulture", "Omnivore", "Meso-carnivore", "Large-carnivore"), " biomass (kg)", sep="")

var.range<-list(c(0, 2, 10, 30),
                c(0, 2, 5, 10),
                c(0, 10, 75, 600),
                c(0, 5, 50, 300),
                c(0, 2, 10, 30),
                c(0, 10, 75, 600))

pdf(file=paste(plot_folder,"FG_boxplots.pdf",sep=""),onefile=T,width=12,height=7)

par(mfrow=c(2,3), cex.lab=1.7, cex.axis=1.5, mar=c(4,5,4,4))

a=0

for (i in sel.var){
  a=a+1
  
    boxplot(dat[, i] ~ landscape, col=col.reg, cex=0.75, , xlab="",ylab=name.var[a], xaxt="n", yaxt = "n")
    
    if (a==1) legend("topleft", c("Spain", "South Africa"), col=c(blues9[4], "orange"), pch=16,  bty="n", cex=2.5,
                     y.intersp=0.75)
   
    mtext(let[a], line = 1.0, adj = -0.2, cex = 1.4, font = 2)
    
    axis(1, 1:5, c("ES", "CZ", "MK", "CC", "HiP"), cex=1.4)
    
    axis(2, log(var.range[[a]]+1), var.range[[a]])
    if(a==5) mtext("Area", line= -22.5, cex=1.5)
  
}
dev.off()


##############################################################################
### Correlations to identify species and biodiversity facets driving carrion consumption
##############################################################################

########## Preparing species data

## Exploratory analysis to select the subset of best predictors 

all.brt<-dat[,c(3, 8:19, 20, 24, 25, 27, 29, 32, 33, 35, 36, 37, 38, 41, 43)+1]

# Type of predictors
pred.type<-c(rep("identity", 7), rep("diversity", 2), rep("equivalence", 3), rep("identity", 13))


### Splitting datasets

ibe.brt<-data.frame(cons_rate=dat[which(dat$region=="ibe"),4], log(taxa[which(dat$region=="ibe"),2:20]+1))
saf.brt<-data.frame(cons_rate=dat[which(dat$region=="saf"),4], log(taxa[which(dat$region=="saf"),21:ncol(taxa)]+1))

ibe.brt2<-data.frame(cons_rate=dat[which(dat$region=="ibe"),4], landscape=dat$sub_region[which(dat$region=="ibe")], log(taxa[which(dat$region=="ibe"),2:20]+1))
saf.brt2<-data.frame(cons_rate=dat[which(dat$region=="saf"),4], landscape=dat$sub_region[which(dat$region=="saf")], log(taxa[which(dat$region=="saf"),21:ncol(taxa)]+1))

ddply(ibe.brt2, .(landscape), function(x) cor(x[,1], x[,3:ncol(ibe.brt2)], method="spearman")) -> cor.imp.ibe2
ddply(saf.brt2, .(landscape), function(x) cor(x[,1], x[,3:ncol(saf.brt2)], method="spearman")) -> cor.imp.saf2

t(cor.imp.ibe2[,-1])->cor.ibe.df
colnames(cor.ibe.df)<-cor.imp.ibe2[,1]

t(cor.imp.saf2[,-1])->cor.saf.df
colnames(cor.saf.df)<-cor.imp.saf2[,1]

cor.ibe.df <- data.frame(f.gr=f.gr[(intersect(rownames(cor.ibe.df), names(f.gr)))], cor.ibe.df)
cor.saf.df <- data.frame(f.gr=f.gr[(intersect(rownames(cor.saf.df), names(f.gr)))], cor.saf.df)

cor.ibe.df2<-data.frame(spp=rownames(cor.ibe.df), f.gr=cor.ibe.df$f.gr, landscape=rep(cor.imp.ibe2[,1], each=nrow(cor.ibe.df)), cor=c(cor.ibe.df$CC, cor.ibe.df$CZ, cor.ibe.df$ES))
cor.saf.df2<-data.frame(spp=rownames(cor.saf.df), f.gr=cor.saf.df$f.gr, landscape=rep(cor.imp.saf2[,1], each=nrow(cor.saf.df)), cor=c(cor.saf.df$HiP, cor.saf.df$Mkhuze))

cor.df<-rbind(cor.ibe.df2, cor.saf.df2)

ddply(ibe.brt2, .(landscape), function(x) apply(x[,3:ncol(ibe.brt2)],2,function(x) length(which(x>0)))) -> occur.ibe
ddply(saf.brt2, .(landscape), function(x) apply(x[,3:ncol(saf.brt2)],2,function(x) length(which(x>0)))) -> occur.saf

write.table(cor.imp.ibe2, paste(plot_folder,"ibe2_spp_var_imp.txt",sep=""),sep="\t")
write.table(cor.imp.saf2, paste(plot_folder,"saf2_spp_var_imp.txt",sep=""),sep="\t")

write.table(occur.ibe, "ocurr_ibe.txt", sep="\t")
write.table(occur.saf, "ocurr_saf.txt", sep="\t")

cor.df$occur<-c(as.vector(t(occur.ibe[,-1])),as.vector(t(occur.saf[,-1])))

#################################################
# Functional group importance
#################################################

# Molten data.frame
cor.df[which(cor.df$occur>4),]->cor.bio2

# Calculating mean and CI correlation
ddply(cor.bio2, .(f.gr), function(x) sd(x$cor)/sqrt(nrow(x))) -> m.se
ddply(cor.bio2, .(f.gr), function(x) mean(x$cor)) -> m.mean

## Dotplot for metric importance
avg<-m.mean$V1
ci1<-m.mean$V1 - qnorm(0.975)*m.se$V1
ci2<-m.mean$V1 + qnorm(0.975)*m.se$V1

names(ci1)<-names(ci2)<-names(avg)<-c("Corvids",  "Large carnivores", "Meso-carnivores","Omnivores","Other raptors", "Vultures")

# sorting at decreasing order
sort(avg)->avg

# sorting in the same order
ci1[names(avg)]->ci1
ci2[names(avg)]->ci2

pdf(file=paste(plot_folder,"FG_cor2.pdf",sep=""),onefile=T,width=8.5,height=7)

fgr.col<-c("#4600FFFF","#CC00FFFF",blues9[5], "gold3","#00FF66FF","#FF0000FF")

dotplot(avg, xlab=expression(Spearman ~ correlation ~ (rho)), xlim=c(min(ci1)-.1, max(ci2)+.1),
        
        par.settings = list(axis.text = list(cex = 2, font=1), 
                            par.xlab.text = list(cex = 2, font=1), 
                            par.ylab.text = list(cex = 2, font=1)),
        panel=function(x,y){
          panel.segments(ci1, as.numeric(y), ci2, as.numeric(y), lty=1, col=fgr.col)
          panel.xyplot(x, y, pch=c(15), cex=2,col=fgr.col)
          panel.abline(v=0, col="black", lty=2)
          
        })
dev.off()

#################################################
# Species importance
#################################################

# Calculating mean and CI correlation
ddply(cor.bio2, .(spp), function(x) sd(x$cor)/sqrt(nrow(x))) -> m.se
ddply(cor.bio2, .(spp), function(x) mean(x$cor)) -> m.mean

which(is.na(m.se$V1)==T)->spp.na

m.se[spp.na,2]<-0

## Dotplot for metric importance
avg<-m.mean$V1
ci1<-m.mean$V1 - qnorm(0.975)*m.se$V1
ci2<-m.mean$V1 + qnorm(0.975)*m.se$V1

names(ci1)<-names(ci2)<-names(avg)<-gsub("_", " ", m.mean$spp)

# sorting at decreasing order
sort(avg)->avg

# sorting in the same order
ci1[names(avg)]->ci1
ci2[names(avg)]->ci2

met_type<-as.factor(c(rep("id", 11), rep("fd", 2), rep("eq", 3)))

f.gr.sel<-as.factor(f.gr[(intersect(m.mean$spp, names(f.gr)))])
names(f.gr.sel) <- gsub("_", " ", names(f.gr.sel))
f.gr.sel[names(avg)]->f.gr.sel

area<-as.factor(c(rep("ibe",19), rep("saf", length(20:ncol(taxa)))))
names(area)<-gsub("_", " ", names(taxa)[-1])

area<-area[(intersect(names(avg), names(area)))]

pdf(file=paste(plot_folder,"spp_cor.pdf",sep=""),onefile=T,width=11,height=13.5)

fgr.col<-c("#4600FFFF","#CC00FFFF",blues9[5], "gold3","#00FF66FF","#FF0000FF")

dotplot(avg, xlab=expression(Spearman ~ correlation ~ (rho)), xlim=c(min(ci1)-.1, max(ci2)+.1),
        
        par.settings = list(axis.text = list(cex = 2.25, font=3), 
                            par.xlab.text = list(cex = 2, font=1), 
                            par.ylab.text = list(cex = 2, font=1)),
        panel=function(x,y){
          panel.segments(ci1, as.numeric(y), ci2, as.numeric(y), lty=1, col=fgr.col[f.gr.sel])
          panel.xyplot(x, y, pch=c(15,17)[area], cex=2.5,col=fgr.col[f.gr.sel])
          panel.abline(v=0, col="black", lty=2)
          
        })
dev.off()

#################################################
# Importance of scavenging community facets
#################################################

ibe.bio<-data.frame(all.brt[which(dat$region=="ibe"),])
saf.bio<-data.frame(all.brt[which(dat$region=="saf"),])

apply(ibe.brt,2,function(x) length(which(x>0)))->occur.ibe
apply(saf.brt,2,function(x) length(which(x>0)))->occur.saf

ibe.brt[,which(occur.ibe>1)]->ibe.brt
saf.brt[,which(occur.saf>1)]->saf.brt


bio.dat<-data.frame(cons_rate=dat$cons_rate, landscape=dat$sub_region, dat[, c(8:19, 20, 24, 25, 27, 29, 32, 33, 35, 36, 37, 38, 41, 43)])

bio.dat<-data.frame(cons_rate=dat$cons_rate, 
                    landscape=dat$sub_region, 
                    dat[, c("FR1", "FR2", "FR3", "FR4", "FR5", "FR6", "range", "scavenger_obligate", "predator_top_pred", 
                            "diet_carnivorous", "social_social","FRic","FDis", "tax.ric", "abun", "bio")])

ddply(bio.dat, .(landscape), function(x) cor(x[,1], x[,3:ncol(bio.dat)], method="spearman")) -> cor.bio

t(cor.bio[,-1])->cor.df
colnames(cor.df)<-cor.bio[,1]
var.names<-c("Corvid biomass", "Other raptor biomass", "Vulture biomass", "Omnivore biomass", "Meso-carnivore biomass", "Large-carnivore biomass", "Home range", "% obligate scavengers", "% top predators", 
                    "% carnivorous", "% social foraging","Functional richness","Functional dispersion", "Species richness", "Abundance", "Biomass")

rownames(cor.df)<-var.names

# Saving results
write.table(cor.df, paste(plot_folder,"cor_metrics.txt",sep=""),sep="\t")

# Molten data.frame for xyplot
cor.bio2<-data.frame(metric=rownames(cor.df), landscape=rep(colnames(cor.df),each=nrow(cor.df)), cor=as.numeric(cor.df))

# Calculating mean and CI correlation
ddply(cor.bio2[which(is.na(cor.bio2$cor)==F), ], .(metric), function(x) sd(x$cor)/sqrt(nrow(x))) -> m.se
ddply(cor.bio2[which(is.na(cor.bio2$cor)==F), ], .(metric), function(x) mean(x$cor)) -> m.mean

## Dotplot for metric importance
avg<-m.mean$V1
ci1<-m.mean$V1 - qnorm(0.975)*m.se$V1
ci2<-m.mean$V1 + qnorm(0.975)*m.se$V1

names(ci1)<-names(ci2)<-names(avg)<-m.mean$metric

# sorting at decreasing order
sort(avg)->avg

# sorting in the same order
ci1[names(avg)]->ci1
ci2[names(avg)]->ci2

met_type<-as.factor(c(rep("id", 11), rep("fd", 2), rep("eq", 3)))
names(met_type)<-var.names
met_type[names(avg)]->met_type

pdf(file=paste(plot_folder,"metric_cor2.pdf",sep=""),onefile=T,width=10,height=8.5)
dotplot(avg, xlab=expression(Spearman ~ correlation ~ (rho)), xlim=c(min(ci1)-.1, max(ci2)+.1),
        
        par.settings = list(axis.text = list(cex = 2, font=1), 
                            par.xlab.text = list(cex = 2, font=1), 
                            par.ylab.text = list(cex = 2, font=1)),
        panel=function(x,y){
                  panel.segments(ci1, as.numeric(y), ci2, as.numeric(y), lty=1, col=c("brown1", "dark green", "#56B4E9")[met_type])
                  panel.xyplot(x, y, pch=c(15,16,17)[met_type], cex=2,col=c("brown1", "dark green", "#56B4E9")[met_type])
                  panel.abline(v=0, col="black", lty=2)
  
})
dev.off()


############ code to plot legends

pdf(file=paste(plot_folder,"legends.pdf",sep=""),onefile=T,width=12,height=4)

par(mfrow=c(1,2), cex.lab=1.7, cex.axis=1.3, mar=c(4,5,4,4))

fgr.col<-c("#4600FFFF","#CC00FFFF",blues9[5], "gold3","#00FF66FF","#FF0000FF")

plot(1, pch="?")

#plotting legend in the graph
legend("right", legend = c("Corvids", "Other raptors", "Vultures", "Omnivores", "Meso-carnivores", "Large-carnivores"),lwd=3, 
       col=fgr.col[c(3,6,1,2,4,5)], bty = "y", cex=1.5, y.intersp = 0.75)


plot(1, pch="?")
#plotting legend in the graph
legend("bottomright", legend = c("identity", "diversity", "equivalence"),pch=c(17,16,15), 
       col=c("#56B4E9", "dark green", "brown1"), bty = "y", cex=1.5, y.intersp = 0.75)

dev.off()
```
