## Supplementary material for "Large home range scavengers support higher rates of carcass removal": Files including R functions and code to reproduce some of our statistical analyses: 3_scav_models.R

```
#################################################################################################################
#                                                                                                               #
# Supplementary Appendix S1                                                                                     #
#                                                                                                               #
# R script to explore biodiversity drivers of carcass removal                                                   # 
#                                                                                                               #
#                                                                                                               #
# Large home range scavengers support higher rates of carcass removal                                           #
#                                                                                                               #
# Gutiérrez-Cánovas, C., Moleón, M., Mateo-Tomás, P., Olea, P.P., Sebastián-González, E.                        #
# Sánchez-Zapata, J.A.                                                                                          #
#                                                                                                               #
# Code written by Cayetano Gutiérrez-Cánovas,                                             #
#                                                                                                               #
#################################################################################################################

# Working directory
setwd("your_folder")
plot_folder<-paste(getwd(),"/plots/",sep="")
assum_folder<-paste(getwd(),"/plots/assum/",sep="")

# Loading libraries
library(car)
library(lattice)
library(usdm)
library(plyr)
library(MuMIn)
library(nlme)

#### Loading data
dat<-read.table("dat_final.txt",h=T,sep="\t")
taxa<-read.table("taxa_ab.txt",h=T,sep="\t")
traits<-read.table("traits.txt",h=T,sep="\t")
f.gr<-read.table("f_gr.txt",h=T,sep="\t")

# Arranging matrices
dat[,1]->rownames(dat)
dat[,-1]->dat

rownames(f.gr)->spp.names

# Arranging functional group matrix
f.gr<-f.gr$x
names(f.gr)<-spp.names

###########################
# Variable transformation
###########################

q<-sapply(dat,class)=="numeric" | sapply(dat,class)=="integer"# selecting quantitative variables

par(mfrow=c(3,3))
for (i in which(q==T)) hist(dat[,i], main=names(dat)[i])
par(mfrow=c(1,1))

log(log(dat$weight_kg))->dat$weight_kg
log(dat$cons_rate)->dat$cons_rate
log(dat$body_mass)->dat$body_mass
hist(car::logit(dat$activity_both/100))

sqrt(dat$FDis)->dat$FDis

log(dat[,c("FR1","FR2","FR3","FR4","FR5","FR6")]+1)->dat[,c("FR1","FR2","FR3","FR4","FR5","FR6")]

log(dat[,c("FR","FRic","tax.ric","abun","bio")])->dat[,c("FR","FRic","tax.ric","abun","bio")]

car::logit(dat[,c("social_social","scavenger_obligate","predator_non_pred","predator_top_pred",
                  "activity_nocturnal","sight_low")])->dat[,c("social_social","scavenger_obligate",
                                                              "predator_non_pred","predator_top_pred","activity_nocturnal","sight_low")]

par(mfrow=c(3,3))
for (i in which(q==T)) hist(dat[,i], main=names(dat)[i])
par(mfrow=c(1,1))

#######################################################################
# Combined influence of landscape and community scavenger facets
########################################################################

# setting colour for plots
col.reg<-c(blues9[4], blues9[4], "orange", blues9[4], "orange")

# Ordering study area codes
landscape<-ordered(dat$type_reg,labels=c("F-M","FO-M","FO-S","FOP-M","FOP-S"), levels=c("F-M","VF-M","VF-S","PVF-M","PVF-S"))

# Checking collinearity

# all predictive variables
View(round(cor(dat[,9:ncol(dat)]),2))

# Only selected subset of predictors
vifstep(dat[,c("weight_kg","range", "predator_top_pred","FDis", "abun", "tax.ric")])
round(as.dist(cor(dat[,c("weight_kg","range", "predator_top_pred","FDis", "abun", "tax.ric")])),2)
range(round(as.dist(cor(dat[,c("weight_kg","range", "predator_top_pred","FDis", "abun", "tax.ric")])),2))

# Creating the modelling dataset
dat_mod<-data.frame(cons_rate=dat$cons_rate, landscape=factor(landscape, ordered=F), scale(dat[,c("weight_kg","range", "predator_top_pred","FDis", "tax.ric", "abun")]))

##############
# Global model
##############

# Global model - needs to check residual structure (heteroscedasticity)
mod0<-gls(cons_rate~weight_kg+range+predator_top_pred+FDis+tax.ric+abun+landscape,data=dat_mod, method="REML")

# Checking residuals on global model
par(mfrow=c(1,2))
plot(fitted(mod0),resid(mod0,"pearson"),main="Residual homocedasticity")
abline(h=0,col="blue")
hist(resid(mod0,"pearson"),main="Residual normality")
par(mfrow=c(1,1))

# Adding a variance structure to account for between-group high variability
vf<-varIdent(form=~ 1 | landscape)
mod1<-gls(cons_rate~range+predator_top_pred+FDis+tax.ric+abun+landscape, weights=vf, data=dat_mod, method="REML")

# Comparing models
AICc(mod0, mod1) # variance structure improves the model and reduce heteroscedasticity

#### Model selection

# Global model with variance structure
mod<-gls(cons_rate~weight_kg+range+predator_top_pred+FDis+tax.ric+abun+landscape, weights=vf, data=dat_mod, method="ML")

# Global model with interaction term
mod_int_range<-gls(cons_rate~weight_kg+range+predator_top_pred+FDis+tax.ric+abun+landscape+range:landscape, weights=vf, data=dat_mod, method="ML")
mod_int_pred<-gls(cons_rate~weight_kg+range+predator_top_pred+FDis+tax.ric+abun+landscape+predator_top_pred:landscape, weights=vf, data=dat_mod, method="ML")
mod_int_fdis<-gls(cons_rate~weight_kg+range+predator_top_pred+FDis+tax.ric+abun+landscape+FDis:landscape, weights=vf, data=dat_mod, method="ML")
mod_int_ric<-gls(cons_rate~weight_kg+range+predator_top_pred+FDis+tax.ric+abun+landscape+tax.ric:landscape, weights=vf, data=dat_mod, method="ML")
mod_int_abun<-gls(cons_rate~weight_kg+range+predator_top_pred+FDis+tax.ric+abun+landscape+abun:landscape, weights=vf, data=dat_mod, method="ML")

AICc(mod, mod_int_range, mod_int_pred, mod_int_fdis, mod_int_ric, mod_int_abun) # checking for interaction between range and AICc

# Global model including variance structure and interaction term
mod<-gls(cons_rate~weight_kg+range+predator_top_pred+FDis+tax.ric+abun+landscape+tax.ric:landscape, weights=vf, data=dat_mod, method="ML")

# Global model summary
summary(mod)
Anova(mod, type=3)

# Checking residuals on global model
par(mfrow=c(1,2))
plot(fitted(mod),resid(mod,"pearson"),main="Residual homocedasticity")
abline(h=0,col="blue")
hist(resid(mod,"pearson"),main="Residual normality")
par(mfrow=c(1,1))

########################
# Multi-model inference
########################

options(na.action = "na.fail") # necessary to run dredge()

# runs all possible models 	and ranks the output according to the AIC, R2 is printed 
mod_d <- dredge (mod, rank = "AICc", extra =  c("R^2")) 

# Saving results
write.table(mod_d[which(cumsum (mod_d$weight) <= 0.95),],paste(plot_folder,"mod_d.txt",sep=""),sep="\t")

mod_set <- get.models (mod_d, subset=cumsum (mod_d$weight) <= 0.95) # subset 

length(mod_set)

# Selecting the model with the lowest AICc
mod.f<-update(mod_set[[1]],method="REML")

summary(mod.f)

########################
# Variance partitioning
########################

var_set<-mod_d[which(is.na(mod_d$abun) & is.na(mod_d$predator_top_pred) & is.na(mod_d$FDis)),]

r.top<-var_set[1,"R^2"]
r.null<-var_set[which(is.na(var_set$weight_kg) & is.na(var_set$range) & is.na(var_set$tax.ric) & is.na(var_set$landscape) & is.na(var_set$`landscape:tax.ric`)),"R^2"]

r.range<-var_set[which(is.na(var_set$weight_kg) & is.na(var_set$range)==F & is.na(var_set$tax.ric) & is.na(var_set$landscape) & is.na(var_set$`landscape:tax.ric`)),"R^2"]
r.ric<-var_set[which(is.na(var_set$weight_kg) & is.na(var_set$range) & is.na(var_set$tax.ric)==F & is.na(var_set$landscape) & is.na(var_set$`landscape:tax.ric`)),"R^2"]
r.weight<-var_set[which(is.na(var_set$weight_kg)==F & is.na(var_set$range) & is.na(var_set$tax.ric) & is.na(var_set$landscape) & is.na(var_set$`landscape:tax.ric`)),"R^2"]

r.ran_ric<-var_set[which(is.na(var_set$weight_kg) & is.na(var_set$range)==F & is.na(var_set$tax.ric)==F & is.na(var_set$landscape) & is.na(var_set$`landscape:tax.ric`)),"R^2"]
r.weight_ric<-var_set[which(is.na(var_set$weight_kg)==F & is.na(var_set$range) & is.na(var_set$tax.ric)==F & is.na(var_set$landscape) & is.na(var_set$`landscape:tax.ric`)),"R^2"]
r.weight_ran<-var_set[which(is.na(var_set$weight_kg)==F & is.na(var_set$range)==F & is.na(var_set$tax.ric) & is.na(var_set$landscape) & is.na(var_set$`landscape:tax.ric`)),"R^2"]

# unique variance

ru.ric<-r.top-r.weight_ran # 0.01
ru.ran<-r.top-r.weight_ric # 17.9
ru.weight<-r.top-r.ran_ric # 6.4

#null r=16.7

# Saving results
write.table(summary (mod.f)$tTable,paste(plot_folder,"res_best.txt", sep=""),sep="\t")

# Checking residuals on global model
par(mfrow=c(1,2))
plot(fitted(mod.f),resid(mod.f,"pearson"),main="Residual homocedasticity")
abline(h=0,col="blue")
hist(resid(mod,"pearson"),main="Residual normality")
par(mfrow=c(1,1))

# Model weights
mod.weights <- Weights(as.numeric(lapply(mod_set, AICc)))

# Updating fitted values using REML
mod_set<-lapply(mod_set, function(x) update(x, method="REML"))

################################################
######### Plotting model results
################################################

lwd.set=3
col.reg<-c(rep("orange", 2), rep("blue",3))
lty.reg<-c(1,2,1,2,3)

pdf(file=paste(plot_folder,"mod_res_both2.pdf",sep=""),onefile=T,width=13,height=6)

par(mfrow=c(1,2), cex.lab=1.8, cex.axis=1.6, mar=c(4,5,4,4))

plot(dat_mod$cons_rate~dat_mod$range, ylab=expression(Consumption ~ rate ~ (kg~ ~ h^{-1})),
     xlab=expression(Home ~ range ~ (km ^{2})), xaxt = "n", yaxt = "n", main="",
     #col=c("black", "dark grey", "black")[dat$type], pch=c(1,16,16)[dat$type])
     col=c("brown1", "dark green", "#56B4E9")[dat$type], pch=c(15,16,17)[dat$type])

mtext("a)", line = 1.0, adj = -0.2, cex = 2, font = 2)

r.m<-mean(dat$range)
r.sd<-sd(dat$range)
x.labs=(c(1,2,3,4)-r.m)/r.sd

#axis(1,x.labs,round(back_st(x.labs,r.m,r.sd),0))

axis(2, log(c(0.01, 0.3, 5, 150)), c(0, 0.3, 2, 150))
axis(1,x.labs,c("< 10", "10 - 100", "100 - 1000", ">1000"))


r.seq<-seq(min(dat_mod$range), max(dat_mod$range), length.out = 1000)

a=0

fit_land<-matrix(NA, length(r.seq), length(unique(landscape)))

for (i in unique(landscape)){
  a=a+1
  
  mod.pred <- data.frame(lapply(mod_set, function(x) { 
    
    predict(x,data.frame(range=r.seq,
                         landscape=i,
                         predator_top_pred=mean(dat_mod$predator_top_pred),
                         FDis=mean(dat_mod$FDis),
                         weight_kg=mean(dat_mod$weight_kg),
                         tax.ric=mean(dat_mod$tax.ric),
                         abun=mean(dat_mod$abun)))
  }))
  
  # mean weighted prediction
  fit_land[,a] <- apply(mod.pred, 1, function(x) sum(x * mod.weights)) 
  
  
  
}

avg.pred <- rowMeans(fit_land)

lines(r.seq,avg.pred,lwd=lwd.set, col="orange")


### Second plot

plot(dat_mod$cons_rate~dat_mod$tax.ric, ylab=expression(Consumption ~ rate ~ (kg~ ~ h^{-1})),
     xlab="Species richness", xaxt = "n", yaxt = "n", main="",
     col=c("brown1", "dark green", "#56B4E9")[dat$type], pch=c(15,16,17)[dat$type])

mtext("b)", line = 1.0, adj = -0.2, cex = 2, font = 2)

r.m<-mean(dat$tax.ric)
r.sd<-sd(dat$tax.ric)
x.labs=(log(c(1:6,8))-r.m)/r.sd

axis(2, log(c(0.01, 0.3, 5, 150)), c(0, 0.3, 2, 150))
axis(1,x.labs,c(1:6,8))


r.seq<-seq(min(dat_mod$tax.ric), max(dat_mod$tax.ric), length.out = 1000)

a=0

fit_land<-matrix(NA, length(r.seq), length(unique(landscape)))

for (i in unique(landscape)){
  a=a+1
  
  mod.pred <- data.frame(lapply(mod_set, function(x) { 
    
    predict(x,data.frame(range=mean(dat_mod$range),
                         landscape=i,
                         predator_top_pred=mean(dat_mod$predator_top_pred),
                         FDis=mean(dat_mod$FDis),
                         weight_kg=mean(dat_mod$weight_kg),
                         tax.ric=r.seq,
                         abun=mean(dat_mod$abun)))
  }))
  
  # mean weighted prediction
  fit_land[,a] <- apply(mod.pred, 1, function(x) sum(x * mod.weights)) 
  
  
  
}

avg.pred <- rowMeans(fit_land)

lines(r.seq,avg.pred,lwd=lwd.set, col="orange")

dev.off()
```
