## Supplementary material for "Large home range scavengers support higher rates of carcass removal": Files including R functions and code to reproduce some of our statistical analyses: readme.txt

Description of the files of the Supplementary Appendix S1

Paper title: Large home range scavengers support higher rates of carcass removal                
Authors: Gutiérrez-Cánovas, C., Moleón, M., Mateo-Tomás, P., Olea, P.P., Sebastián-González, E., Sánchez-Zapata, J.A.                                                               
Code written by Cayetano Gutiérrez-Cánovas,                  

0_FD_functions.R
R script to generate trait-based metrics

0_quality_funct_space.R
R function for computing the quality of functional multidimensional functional spaces

1_biometric estimation.R
R script to estimate scavenger community metrics (functional identity, functional diversity, functional equivalence)

2_descriptive_plots.R
R script to create descriptive plots and figures

3_scav_models.R
R script to explore biodiversity drivers of carcass removal

traits.txt
Scavenger traits
